# Hybrid molecular dynamics–deep generative framework expands *apo* RNA ensembles toward cryptic ligand-binding conformations: application to HIV-1 TAR

**DOI:** 10.1101/2025.01.07.631832

**Authors:** Ikuo Kurisaki, Michiaki Hamada

## Abstract

RNA plays vital roles in diverse biological processes and represents an attractive class of therapeutic targets. In particular, cryptic ligand-binding sites—absent in *apo* structures but formed upon conformational rearrangement—offer high specificity for RNA–ligand recognition, yet remain rare among experimentally-resolved RNA–ligand complex structures and difficult to predict *in silico*. RNA-targeted structure-based drug design (SBDD) is therefore limited by challenges in sampling cryptic states. Here, we apply Molearn, a hybrid molecular dynamics–deep generative framework, to expand *apo* RNA conformational ensembles toward cryptic states. Focusing on the paradigmatic HIV-1 TAR–MV2003 system, Molearn was trained exclusively on *apo* TAR conformations and used to generate a diverse ensemble of TAR structures. Candidate cryptic MV2003-binding conformations were subsequently identified using post-generation geometric analyses. Docking simulations of these conformations with MV2003 yielded binding poses with RNA–ligand interaction scores comparable to those of NMR-derived complexes. Notably, this work provides the first demonstration that a generative model can access cryptic RNA conformations that are ligand-binding competent and have not been recovered in prior molecular dynamics and deep generative modeling studies. Finally, we discuss current limitations in scalability and systematic detection, including application to the Internal Ribosome Entry Site, and outline future directions toward RNA-targeted SBDD.

## Introduction

RNA molecules are involved in a wide variety of biological processes ^1–4^. Their dysfunctions bring about serious diseases and they have been regarded as important targets for drug discovery ^5–8^. Recalling that Structure-Based Drug Design (SBDD) has drawn much attention for the monumental successes in protein-targeted drug discovery ^9–13^, utilizing tertiary structures of RNA is similarly a promising approach to systematically discover RNA-targeted drugs among abundant candidate compounds ^14^. In the line of SBDD approach, if we could design drugs interacting with a cryptic RNA–ligand binding site, which only emerges in an RNA–ligand complex, such drugs should be underscored due to higher specificity to target RNA and weaker side effects for off-target RNA molecules. Meanwhile, although the number of RNA–ligand complex structures has increased, it remains insufficient, thus hindering development of RNA-targeting SBDD. As of January 27, 2026, experimentally derived, protein-free RNA–ligand complex structures comprise approximately 600 entries in the Protein Data Bank (PDB), corresponding to about 2.5% of all PDB entries, as cataloged by the Nucleic Acid Knowledgebase ^15^. RNAs harboring cryptic binding sites are even more sparsely characterized. It therefore remains infeasible to routinely perform cryptic binding site-targeted SBDD for arbitrary RNA targets.

Because of the paucity of experimentally resolved RNA structures, theoretical prediction of *apo* RNA harboring cryptic binding site is expected as the initial step to generate RNA–ligand docked poses. However, theoretical approaches have not achieved such tasks so far, even for a paradigmatic RNA to study interaction with ligands, HIV-1 TAR (Human Immunodeficiency Virus type 1 Trans-Activation Response element, referred to as TAR hereafter; **Fig. 1A** and **1B** for the second and tertiary structures, respectively)^16–23^. Among TAR-binding ligands, MV2003 (Arginine 4-methoxy-β-napthylamide) is known as a small molecule to bind to TAR’s cryptic binding cavity ^21^ (**Fig. 1C** and **1D** for MV2003-bound and *apo* TAR conformations, respectively). There have been several molecular dynamics (MD) simulation studies to try generating TAR conformations competent for MV2003 binding (referred to as ‘MV2003-bindable TAR conformations’ hereafter) from the *apo* forms ^24–26^. However, none of these MD studies succeeded in generation of MV2003-bindable TAR conformations.

**Figure 1.**
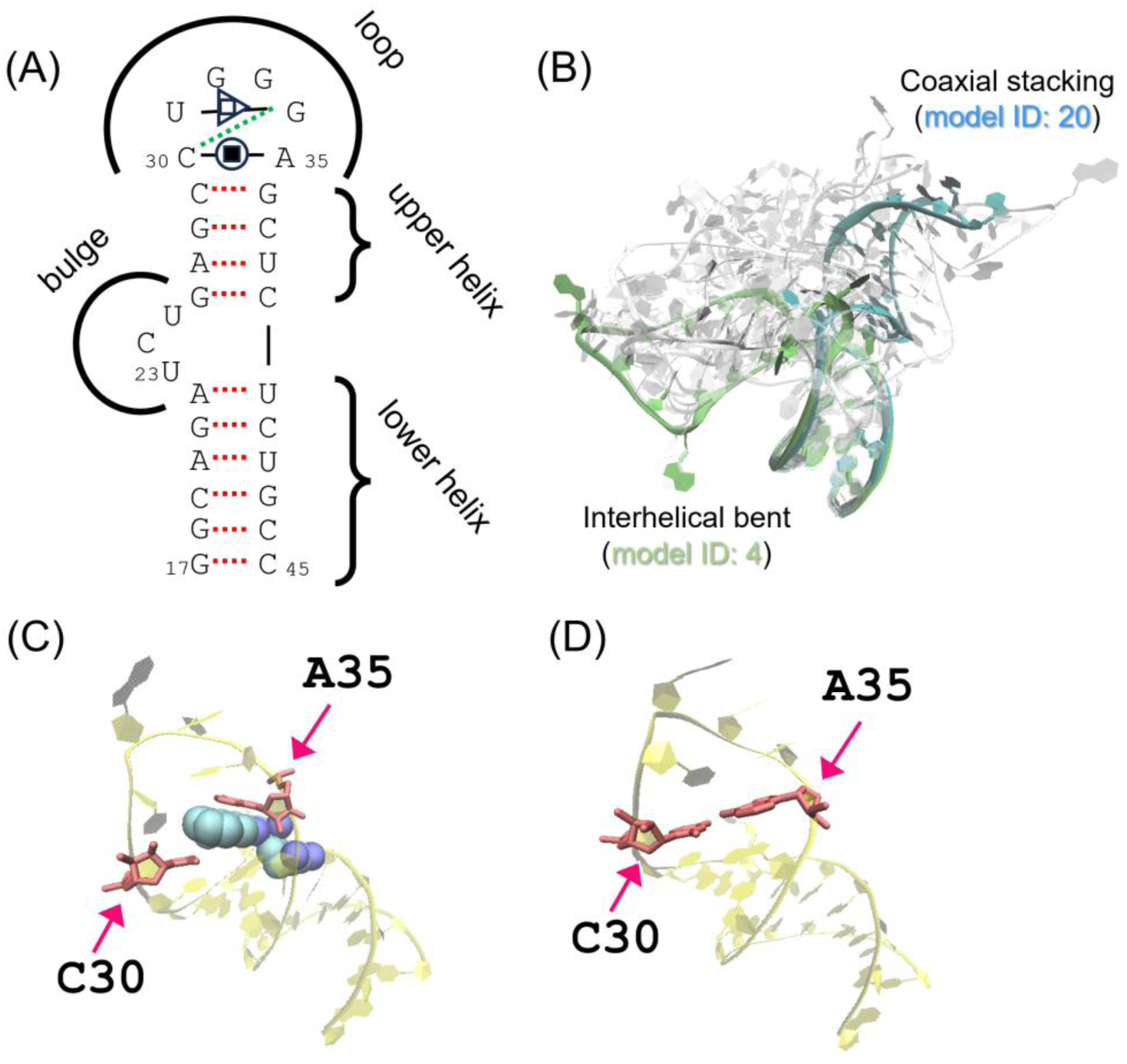
Structures of HIV-1 TAR. (A) Secondary structures and (B) Tertiary structure ensemble in *apo* form (PDB entry: 7JU1), where all of twelve TAR models are illustrated. (C) MV2003-bound structure (model No. 1 of PDB entry: 2L8H). (D) MV2003-inaccessible structure (model No. 1 of PDB entry: 7JU1). In panel A, models 4 and 20, a pair with the largest RMSD value of the all pairs (_20_*C*_19_ = 190) is simultaneously considered. Dotted lines are for Watson-Crick base pair (WC). WC in green only forms in the model No. 4. Closed square in circle and open square in triangle are for non-WC interactions. In panel B, models 4 and 20 are highlighted by green and blue, respectively. In panels C and D, MV2003 and the interacting residues are shown by van der Waals spheres and red sticks, respectively. TAR structure is illustrated by transparent yellow ribbon.

Such negative results likely reflect the difficulty of sampling rare, collective conformational rearrangements within feasible MD timescales, rather than the complete absence of relevant local fluctuations. Indeed, MD simulations with fixed-charge (additive) force-field parameters remain widely used for RNA conformational sampling ^24–26^ owing to their strong track record in reproducing selected physicochemical observables ^27–30^, favorable computational cost, and the still-limited maturity of polarizable force fields developed to improve electronic interaction accuracy ^31^. At the same time, it is well appreciated that fixed-charge RNA force fields can exhibit imbalances in electrostatic interactions, including between base stacking and base-pair hydrogen bonding ^28, 32^. In the case of TAR, stacking interactions near C30 and A35 may be stabilized relative to disruption events required for cryptic cavity formation (**Fig. 1A**, **1C**, and **1D**), further reducing the likelihood that such states are sampled spontaneously in conventional MD. Importantly, however, the absence of cryptic-pocket conformations in available MD trajectories does not imply that the underlying conformational degrees of freedom are inaccessible, but rather that their collective activation is rare. Consistent with previous observations that long or enhanced MD simulations can populate non-native states depending on the force field ^33^, MD alone may not yet constitute a general standalone route to cryptic-site discovery, and is likely to be most effective when combined with complementary approaches.

To overcome the inherent limitations of straightforward MD simulations, we employ a hybrid MD–deep generative framework, Molearn^34^. Importantly, in this context, MD simulations are not intended to provide complete transition pathways or fully accurate energetics, but rather to supply dense and internally consistent local conformational fluctuations for model training. Degiacomi and colleagues proposed a convolutional neural network (CNN) model to systematically generate a set of intermediate protein conformations along the transition path, capturing the global structural reconfigurations that connect the two endpoint states (**Fig. 2** shows a technical outline; detailed explanation is given in **Materials and Methods**). Notably, the earlier Molearn studies succeeded in generating biologically relevant intermediate conformations of proteins and reported performance superior to conventional methods in terms of experimentally resolved transition pathways ^34, 35^.

**Figure 2.**
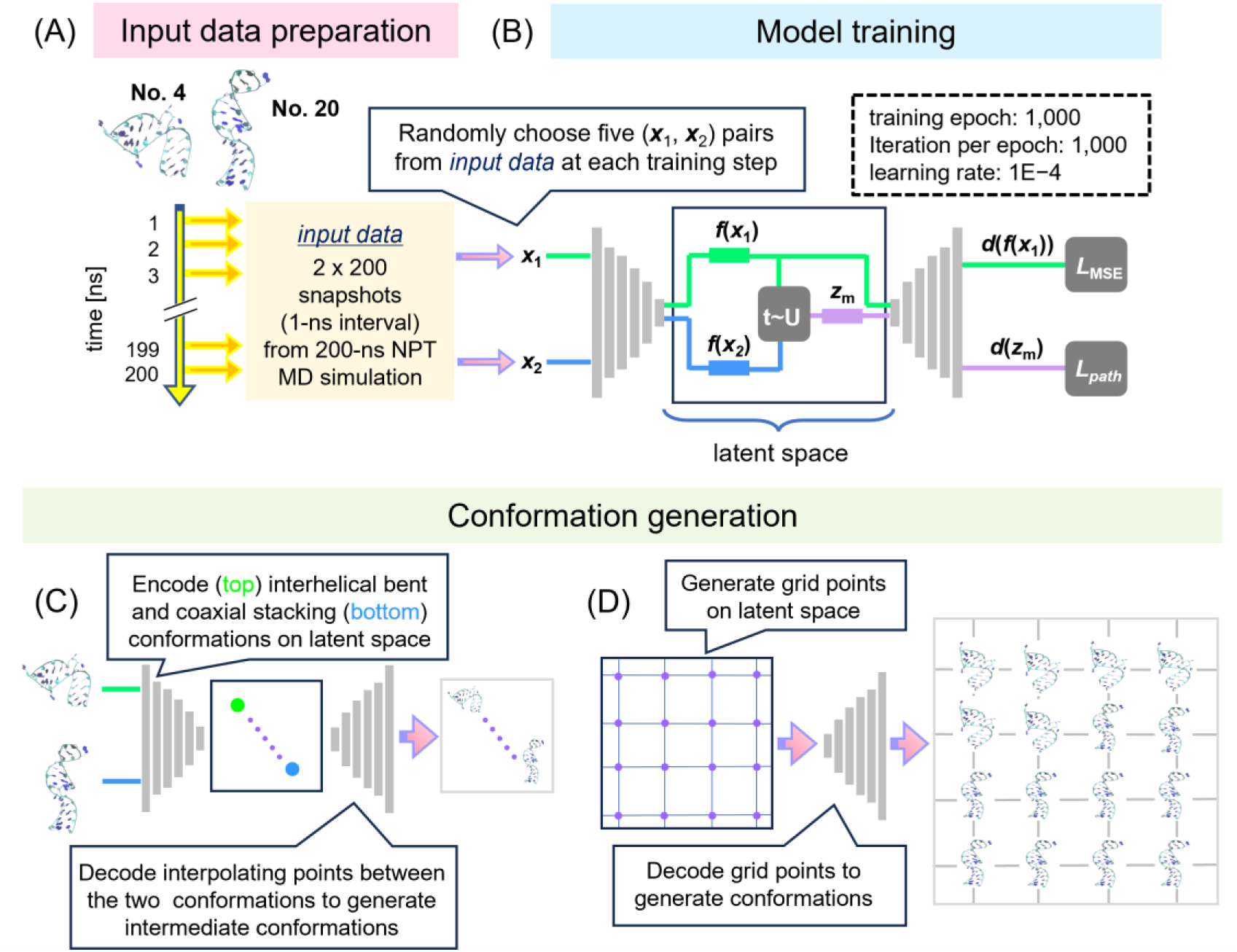
Procedure of Molearn model training and conformation generation. (A) Preparation of input dataset with MD simulations for NMR models No. 4 and No. 20 (PDB entry: 7JU1). (B) Molearn model training. (C) Conformation generation by interpolating two representative conformations, a pair of MD snapshot structures with the maximum RMSD value, on the two-dimensional latent space. (D) Conformation generation from grid points on the latent space. In panel B, gray rectangles denote one dimensional convolution layers. Atomic coordinates are represented by **x_1_** and **x_2_**. Symbols *f* and *d* are for encoded and decoded atomic coordinates. *L*_MSE_ and *L*_path_ are mean square error-based and physics-based loss functions (*see* Materials and Methods for definitions). Latent space coordinate, z_m_, is interpolation point defined by *tf*(***x***_1_) + (1 − *t*)*f*(***x***_2_), where *t* is a value randomly taken from [0, 1] shown as **U** in the panel.

From practical point of view, Molearn’s CNN models (referred to as Molearn models, hereafter) are trained with MD-derived conformational ensembles at equilibrium states, and transition pathways are internally defined on a two-dimensional latent space (*see* **Fig. 2A** and **Fig. 2B**). These features could make Molearn a promising approach to systematically generate ligand-bindable *apo* RNA conformations at a realistic computational cost. A ligand-bindable RNA is expected to exhibit partial disruption of canonical RNA interactions such as base-pairing or base-stacking, due to forming intermolecular interactions with ligands. Such intra-RNA interactions are often disrupted through global conformational changes along transition pathways, rather than within statistically plentiful conformations captured by experimental methods such as X-ray crystallography. Generally, such conformational changes progress within time period of several milliseconds or longer ^36–39^ (as for TAR, disruption of hydrogen bonds formed between C30 and A36 followed by rearrangement of the stem loop), thus being inaccessible even for maximally-long non-equilibrium MD simulations of today. Using Molearn, we could obtain transition pathways undergoing global conformational changes with routinely-performed equilibrium MD simulations reachable within realistic computational time.

In this study, we trained Molearn models on MD-derived *apo* TAR conformations and sought to generate MV2003-bindable TAR conformations. The original Molearn study considered protein backbone atoms for conformation generations ^34^. Meanwhile, we trained Molearn with full-atom RNA models (27 heavy atoms per RNA residue, comprising phosphate, sugar and base) with suitable modification of the distributed Molearn programs. Full-atom RNA representation enables us to quantitatively test ligand-bindable RNA conformations by employing an RNA–ligand docking simulation and an RNA–ligand docking score estimation. In addition, we assessed scalability by applying this hybrid MD–deep generative approach to a larger RNA molecule. Together, these results provide an initial step toward a practical framework to generate RNA conformations with cryptic ligand-binding sites without prior complex structures.

## Materials and methods

### System setup

We considered the NMR-derived *apo* TAR structures (PDB entry ID: 7JU1) as the conformation ensemble in the ground state ^40^ (*see* **SI-1** and **Fig. S1** in Supporting Information for comparison with other two *apo* structure ensembles). Among the 20 conformations, the models No. 4 and No. 20 are selected as the representative interhelical bent and the coaxial stacking conformations, respectively (*see* **Fig. 1B**) because they are the most structurally different pair among the 190 pairs (*i.e.*, _20_C_2_) in terms of Root Mean Square deviation (RMSD).

Each of the two *apo* TAR structures are solvated in the rectangular box with 22580 water molecules, and electronically neutralized with 28 Na^+^. Forces acting among atoms were calculated using AMBER ff99bsc0χOL3 force field ^41^, TIP3P water model ^42, 43^, and Joung/Cheatham ion parameters parameterized for the TIP3P water model ^44, 45^, applied to RNA residues, water molecules, and ions, respectively. In this study, the MD simulations were performed within a relatively short timescale compared with reported timescales of large-scale RNA conformational transitions ^46–49^, and were intended to sample conformations around predefined, thermodynamically stable states−namely, interhelical bent and coaxial stacking states (*see* **Fig. 1B**). Accordingly, we treated the choice of force-field parameters as a modeling assumption, under the premise that it would not qualitatively affect the structural diversity of the sampled conformational ensemble required for Molearn training.

### Molecular Dynamics Simulations

First, atomic steric hindrances were removed using molecular mechanics (MM) simulations, which consist of 1000 steps of steepest descent method followed by 49000 steps of conjugate gradient method. Then, using molecular dynamics (MD) simulations, the system temperature and pressure were relaxed around 300 K and 1 bar, respectively. The following 200-ns NPT MD simulations (300 K, 1 bar) were performed to prepare the training set for Molearn training (*cf.* **Fig. 2A**).

Each MD simulation was performed under the periodic boundary condition. Electrostatic interaction was treated by the Particle Mesh Ewald method, where the real space cutoff was set to 9 Å. The vibrational motions associated with hydrogen atoms were frozen by the SHAKE algorithm. The translational center-of-mass motion of the whole system was removed by every 500 steps to keep the whole system around the origin, avoiding the overflow of coordinate information from the AMBER MD trajectory format. The system temperature and pressure were regulated with Berendsen thermostat ^50^ with a 5-ps coupling constant and Monte Carlo barostat with volume change by each 100 steps, respectively. A set of initial atomic velocities were randomly assigned from the Maxwellian distribution at 0.001 K at the beginning of an MD simulation. The time step of integration was set to 2 fs. Each of MD simulations was performed using the AMBER 22 ^51^ GPU version PMEMD module based on SPFP algorithm ^52^ with NVIDIA GeForce RTX 3090. Further details for simulation procedures are given in **SI-2** in Supporting Information.

### Molearn model and the training procedure

We obtained the Molearn program suite from GitHub in October 2022 (https://github.com/Degiacomi-Lab/molearn/tree/diffusion) and partially modified specific modules to support RNA atom definitions and the corresponding force-field parameter files ^41^. The modified components are distributed as molearnA (*see* Code availability for the repository information). The Molearn model for this RNA study consists of twelve layers (six encoder and six decoder layers) (**Fig. 2B**) ^34^. The first layer has 3 input channels and 32 output channels and the subsequent layers with depth *t* have ⌊32 × 1.8^*t*^⌋ input where batch normalization and rectified linear unit activations are considered.

The training dataset is made from the 200-ns NPT MD trajectories by removing water and ion atoms from snapshot structures. Each snapshot structure is superposed on the NMR model No. 1 of TAR (PDB entry: 7JU1) using root-mean-square fitting algorithm with phosphate and C4’ atoms in the stem region (residues number are from 17 to 21 and from 41 to 45, *see* **Fig. 1A**). This treatment is indispensable due to that a Molearn model uses the Cartesian coordinates as input information so far and is not equivalent/invariant to translation and rotation ^34^. The Molearn model employs size 4 kernels with strides of 2 and padding of 1 in the convolution process. The total number of trainable network parameters is 5414899, which occupies disk space of 445 MB.

We considered all non-hydrogen atoms in RNA residue (27 atom types in total) to train a Molearn model. It is possible to execute the current version of Molearn program with fewer atom sets (linear, non-cyclic backbone trace, and minimum sidechain atom for coarse grained representation of nucleic acid bases; P, O3’, O5’, C4’, C3’, C5’, O4’, C1’, N1, N9, C2, C4), while we here used a full set of non-hydrogen atoms to generate full atom RNA structures.

We consider the following loss function to evaluate training processes

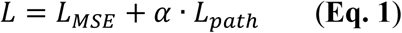

*L_MSE_* is a mean square error (MSE) calculated for sets of atomic coordinates of the original and reconstructed snapshot structures (corresponding to ***x*_1_** and *d*(*f*(***x*_1_**)) in **Fig. 2B**, respectively). *L_path_* is atomistic potential energies for intermediate structures generated by Molearn (represented by ***d*(z_m_)** in **Fig. 2B**). The energy terms consist of atomic bonding, atomic angle, dihedral angle, and approximated non-bonded interaction defined by Coulomb interactions and 12 repulsive Lennard-Jones potential ^34^. Force-field parameters are derived from AMBER ff99bsc0χOL3 force field ^41^.

At every optimization step, the scaling factor *α* is dynamically given by benchmarking an MSE value as follows. *L_path_* is repeatedly multiplied by 0.1 until *L_path_* becomes smaller than *L_MSE_*. This dynamic assignment automatically adjusts the strength of *L_path_* to *L_MSE_* through all training period.

Adam algorithm was employed to optimize Molearn’s parameter set. The model training procedure consists of 1000 epochs, and one epoch has 1000 optimization iteration steps. The learning rate is set to 1e^−4^ and the weight decay is set to 0 as does the distributed Molearn running script. In each optimization step, 5 pairs of snapshot structures are randomly selected from the training dataset as a batch data. For each pair, either of the two is used to calculate *L_MSE_*, while *L_path_* is calculated from a set of Molearn-generated intermediate structures. A midpoint of the pair on the Molearn’s two-dimensional latent space is randomly selected and then decoded into a tertiary structure.

Molearn models were trained using 5 nodes on CPU machines (AMD E7763) of Research Center for Computational Science (RCCS) and took around 6 days for a single 1000 epoch training.

Each of Molearn-generated RNA structures was energetically relaxed using QRNAS program ^53^. An optimization step was repeated by 20000 times with non-bonded interaction cutoff of 12Å under the implicit solvation model. A QRNAS computation for TAR takes about 36 minutes with single core of the RCCS’s CPU machines. Since we finally obtain 10201 TAR conformations from each trained model (meaning of the number will be explained in Results and Discussion section), completing all 10201 QRNAS computations take 12 hours with 500 parallel computations in RCCS.

### Structure analyses

Analyses of RMSD and calculation of geometrical properties (bond length and bond angle involved in RNA) were performed using Cpptraj module of AMBER22 ^51^. We evaluate chemical accuracy of atomic RNA structures using %*_TotalErr_*.

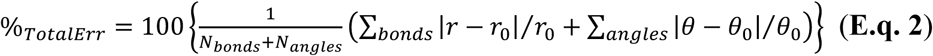

This is an average of deviation of atomic bond length (*r*) and bond angle (*θ*) from their equilibrium values (*r*_0_ and *θ*_0_, respectively). *N_bonds_* and *N_angles_* denote the numbers of bonds and angle values, respectively.

RNA BRiQ scores ^54^ were calculated for experimentally resolved and Molearn-generated RNA conformations as a knowledge-based measure of deviation from canonical RNA geometries. POVME2 is used to calculate atomic volume of cavity in the vicinity of C30 and A35 ^55, 56^. Molecular structures are illustrated using Visual Molecular Dynamics (VMD) ^57^. RNA–ligand docking simulations are performed using RLDOCK ^58^, after which docking poses with anomalous steric clashes are filtered using PoseBusters ^59^, and the remaining poses are evaluated for canonical RNA–ligand interaction patterns using the AnnapuRNA scoring function ^60^. Technical details for POMVE2, RLDOCK and PoseBusters are given in **SI-3**, **SI-4**, and **SI-5** in Supporting Information, respectively. In-house Python scripts for data analysis were prepared with assistance from ChatGPT-4o (OpenAI) and were reviewed and validated by the authors.

## Results and discussion

### Structural descriptors for identifying MV2003-binding cavities in TAR

We examined two types of descriptors for NMR-derived TAR conformations, (i) an RNA-specific knowledge-based score and (ii) geometrical conditions associated with formation of the MV2003-binding cavities. These descriptors provide essential structural context for the present analysis and have not been explicitly characterized in previous studies. ^21^

Using the knowledge-based score, we asserted whether TAR conformations in the MV2003-bound state deviates from canonical RNA geometries observed in the *apo* ensemble. In this case study, formation of the MV2003-binding cavity requires partial disruption of canonical RNA interactions. To evaluate such conformational deformation, we calculated the state-of-the-art RNA-specific knowledge-based score, RNA BRiQ ^54^. The BRiQ scores of TAR in the MV2003-bound state (red squares in **Fig. 3A**) are consistently higher than those of the *apo* conformations (green squares in **Fig. 3A**). These observations indicate that the TAR adopts the MV2003-bound state through disruption of canonical RNA interactions, thereby supporting our strategy of generating MV2003-bindable TAR conformations using Molearn.

**Figure 3.**
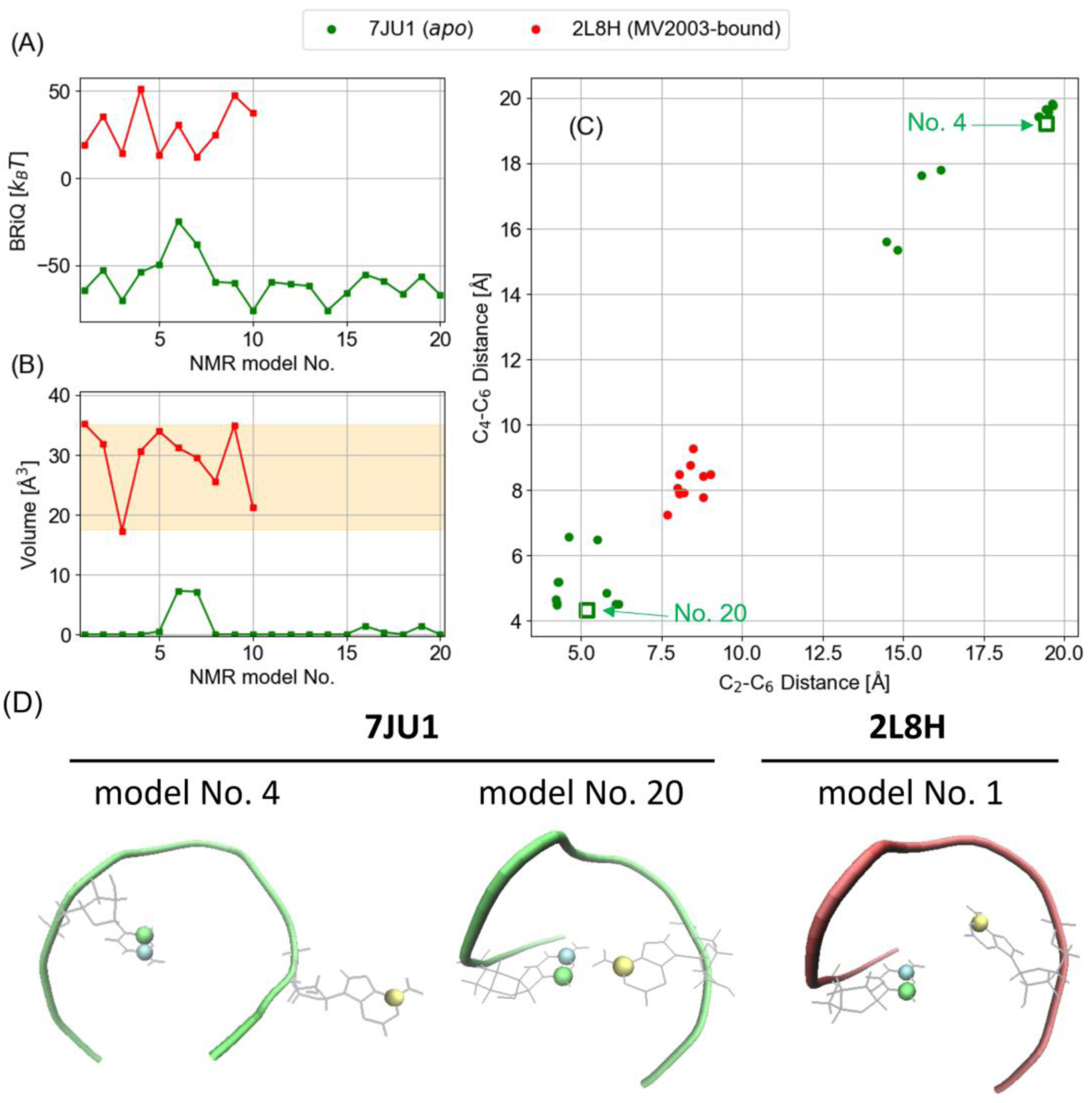
Structural characterization of NMR-derived HIV-1 TAR conformations (*apo* and MV2003-bound forms annotated by PDB entries of 7JU1 and 2L8H, respectively). (A) BRiQ score (BRiQ). (B) Volume of (potential) MV2003-binding cavity. (C) Interatomic distances characterizing positional relationship between C30 and A35. (D) Location of C30 and A35 in *apo* and MV2003-bound forms. In panels A and B, orange belts denote value ranges for volume size of MV2003-binding cavity and BRiQ for TAR conformations in the MV2003-bound state. In panels C, green open circles at top-right and bottom-left are for interhelical bent and coaxial stacking conformations (*c.f.*, **Fig. 2B**), respectively. In panel D, blue, green and yellow balls represent C30’s C_2_, C30’s C_4_ and A35’s C_6_ atoms, respectively.

Next, we consider geometrical conditions satisfied by TAR conformations in the MV2003-bound state. To clearly distinguish geometrical compatibility from experimentally validated binding, throughout this study, we use the term ‘binding-compatible’ to denote conformations that satisfy geometrical requirements for ligand binding. In the following sections, we will systematically generate TAR conformations with trained Molearn models and then comprehensively search MV2003-binding-compatible ones, which have a cavity accessible for MV2003. However, there have been no earlier studies to quantitatively characterize the cavity with geometrical metrics, to our best knowledge. Aiming to perform comprehensive search for TAR conformations of interest, we define geometrical conditions to form the MV2003-binding cavities.

We analyzed volume of cavity formed between the Cyt30 and Ade35 for each of *apo* and MV2003-bound TAR conformations (**Fig. 3B**). Each of MV2003-bound TAR conformations has an MV2003-binding cavity, and the values of cavity volume range from 17 to 35 Å^3^ (*see* red squares in **Fig. 3B**, where the value range is highlighted by the orange belt). Meanwhile, the *apo* TAR conformations do not form sufficiently large cavities (*see* green squares in **Fig. 3B**). The NMR models No. 6 and No 7 of *apo* TAR exhibit relatively large volume values, although these remain around 8 Å^3^ and do not reach the range required for MV2003 binding. The BRiQ scores for these NMR models are relatively higher than those of other *apo* conformations (*see* **Fig 3A**), suggesting small deviation from canonical intra-RNA interactions.

A relative position between MV2003-interacting RNA bases is also an important metric for the cryptic MV2003-binding cavity. The disruption of the RNA base pair results in the formation of the MV2003-binding cavity (**Fig. 1C** and **1D**). The methoxy naphthalene group of MV2003 is bound to TAR by a pseudo-stacking with Cys30 and Ade35, which are found below the apical loop (*see* **Fig. 1C** for the tertiary MV2003-bound structure). Then, we characterized a positional relationship between these RNA bases using a pair of inter-atomic distances. One is distance between C_2_ of Cyt30 and C_6_ of Ade35, and the other is that between C_4_ of Cyt30 and C_6_ of Ade35 (hereafter, the former and latter are referred to as d_2-6_ and d_4-6_, respectively). We can find that the (d_2-6_, d_4-6_) distribution for *apo* conformations and that for MV2003-bound conformations is mutually exclusive (**Fig. 3C**). The MV2003-bound conformations are localized in the (d_2-6_, d_4-6_) space and the distribution is unimodal (a representative case is illustrated in the right panel in **Fig. 3D**). The value ranges of d_2-6_ and d_4-6_ are [7.7, 9.0] and [7.2, 9.3] (Unit: Å), respectively, which is a necessary condition for TAR to form pseudo-stacking with MV2003.

Meanwhile, the distribution for *apo* TAR conformations is bimodal and is located outside of that for the MV2003-bound TAR conformations. These observations denote that any of the NMR-derived *apo* TAR conformations does not take MV2003-binding-compatible configurations for Cyt30 and Ade35. For example, the representative interhelical bent and coaxial stacking conformations (the NMR models No. 4 and No. 20 in **Fig 1B**, highlighted by open green squares in **Fig 3C**) are different in positional relationship of the RNA bases. Ade35 is flipped out in the model No. 4, while Cys30 and Ade35 make contacts in the model No. 20 (*see* the left and center panels in **Fig. 3D**). These arrangements of the bases inhibit pseud-stacking formation with a methoxy naphthalene group of MV2003.

According to the above geometrical analyses, predicted MV2003-binding-compatible conformations should satisfy the three geometrical conditions at the same time: (i) the cavity volume ranges from 17.5 to 35 Å^3^; (ii) d_2-6_ is found between 7.7 and 9.0 Å; (iii) d_4-6_ is found between 7.2 and 9.3 Å. These geometric criteria, derived exclusively from experimentally resolved TAR–MV2003 complexes, were applied post hoc as an evaluation filter and were not imposed as constraints during Molearn training or conformational sampling. Accordingly, ligand-binding compatibility is evaluated independently of the generative model.

### Generative expansion of *apo* TAR conformations toward MV2003-binding-compatible cavity geometries

As in the case of the original Molearn study, we trained Molearn models using a set of snapshot structures obtained from atomistic MD simulations. We performed the two 200-ns NPT MD simulations for TAR. One and the other start from the NMR-derived TAR models No. 4 and No. 20 of 7JU1 (**Fig. 1B**), respectively. Each of the two MD trajectories exhibits equilibrated regions corresponding to representative TAR conformational states: the interhelical bend and coaxial stacking conformations (*see* **SI-2** and **Fig S2** for related discussion). We then picked up 200 snapshot structures from each of the MD trajectories with 1-ns interval. These two sets of 200 snapshot structures are integrated to use as the training dataset (*cf.* **Fig. 2A**).

Notably, none of the 400 MD-derived TAR conformations exhibits a potential MV2003-binding cavity. These conformations show markedly smaller cavity volume of cavity between Cyt30 and Ade35 (green and blue graphs in **Fig. 4A**) compared with the NMR-derived TAR conformations in the MV2003-bound state (the value range for such conformations is given orange belt in **Fig. 4A**). Although the corresponding (d_2-6_, d_4-6_) distributions of the MD-derived conformations (green and blue squares in **Fig. 4B**) are spatially expanded by the influence of thermal fluctuation, they remain distinct from those of the NMR-derived MV2003-bound TAR conformations (red squares in **Fig. 4B**). At the same time, the broadened distributions in Fig. 4B indicate the presence of appreciable local conformational fluctuations in the MD ensemble, even in the absence of explicit cryptic-pocket formation. In the context of generative modeling, such continuous and correlated structural variability provides a practical basis for learning interpolative geometric variations beyond those directly sampled in MD trajectories. Accordingly, Molearn was trained exclusively on MD-derived *apo* TAR conformations that do not exhibit cryptic MV2003-binding cavities, and no structural information from MV2003-bound TAR conformations was used during model training.

**Figure 4.**
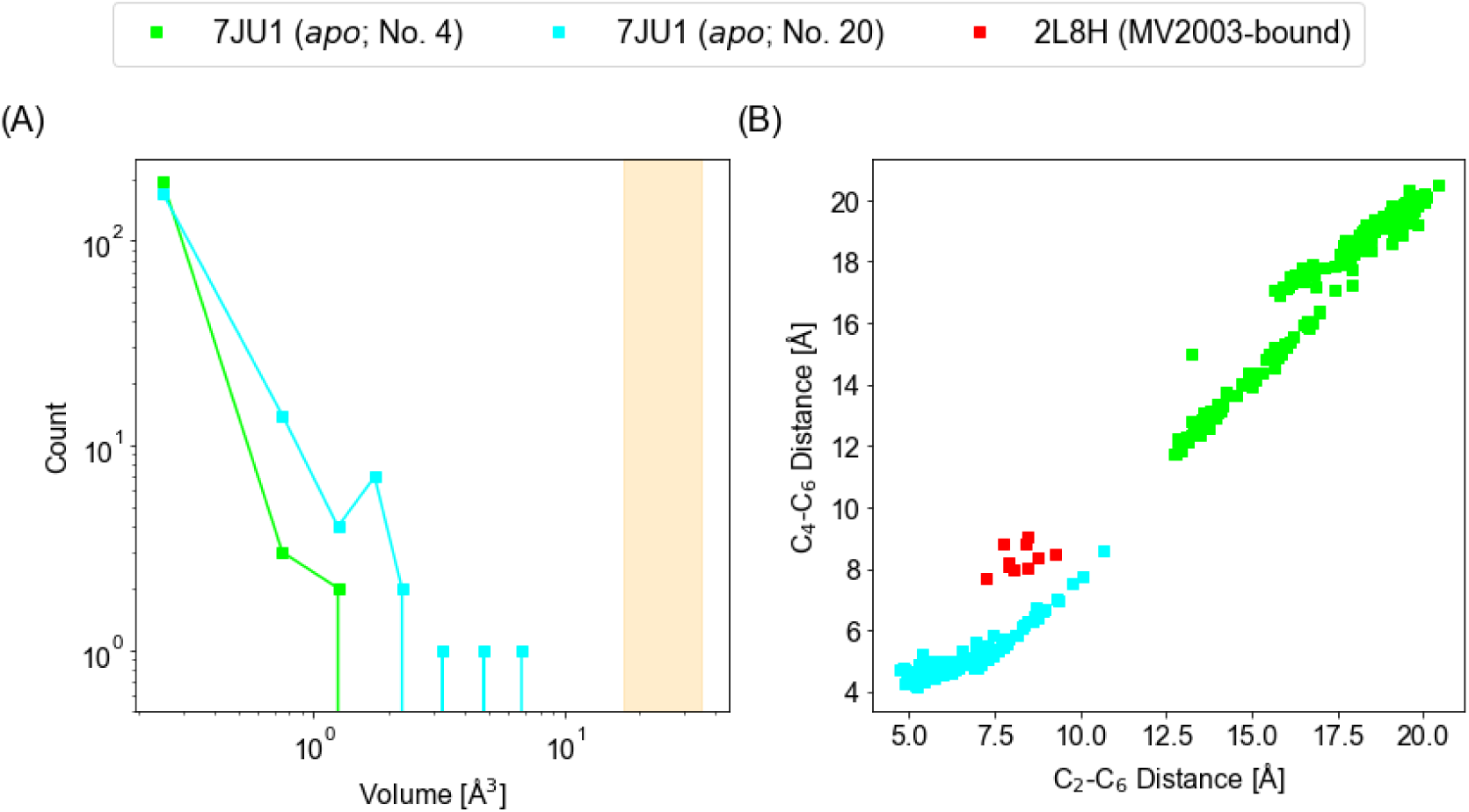
Geometrical properties of MD derived-Molearn input data. (A) Volume of (potential) MV2003-binding cavity. (B) A pair of interatomic distances characterizing positional relationship between C30 and A35. Green and blue squares are for snapshot structures obtained from MD simulations starting from *apo* HIV-1 TAR conformations annotated by No. 4 and No. 20 (PDB entry: 7JU1). In panel A, orange belt denotes value ranges for volume of MV2003-binding cavity for NMR-derived TAR conformations in the MV2003-bound state (PDB entry: 2L8H). In panel B, red squares are for the NMR derived conformations.

Using the training dataset, we independently trained Molearn models ten times. Each of trained Molearn models shows decreasing network-loss through the training epochs and the values appear to converge by 1000^th^ epoch (**Fig. 5A**). We assume that each of Molearn models are finely trained to reproduce the training data and to generate latent space-interpolated intermediate structures with chemically reasonable geometry (*cf.* Molearn’s loss function defined in **Eq. 1**).

**Figure 5.**
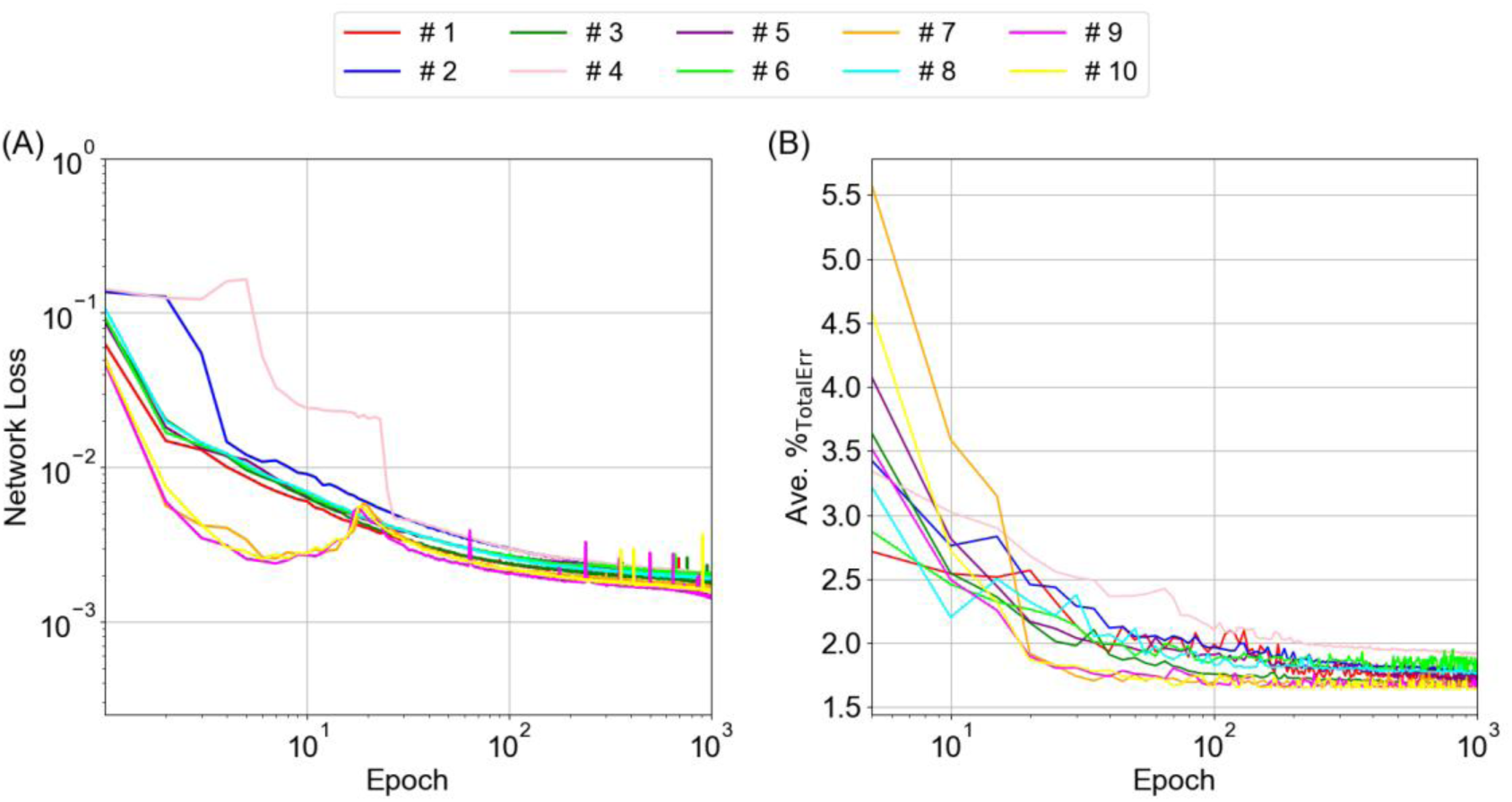
Evaluation of trained Molearn model for HIV-1 TAR. (A) Network loss along training epoch. Value at each epoch is an average over 1000 training steps in the epoch. (B) Atomic precision of Molearn-generated conformations along training epoch. #1 to #10 in the legend box denote the ten independently trained Molearn models. With regard to (B), %_TotalErr_ values are calculated for 50 TAR conformations generated by interpolating the interhelical bent and coaxial stacking forms on Molearn’s latent space (*c.f*. **Fig. 1C** and **Fig. 2B**) at each epoch. A set of 50 %_TotalErr_ values are averaged at each epoch to show the graph.

We confirmed the assumption by calculating the chemical accuracy of Molearn-generated intermediate conformations with %_TotalErr_ (**Eq. 2**). Intermediate structures were generated by interpolating the representative conformations (*cf.* **Fig. 2C**), the interhelical bent and coaxial stacking ones (a pair with the maximum RMSD value picked up from the training dataset, which conformationally resembles the NMR models No. 4 and No. 20 of 7JU1 in **Fig 2B**, respectively) on the two-dimensional latent space learned by each Molearn model. We used 48 latent space coordinates connecting between the two input conformations to generate intermediate TAR structures. They are combined with the two representative conformations, which are used as the terminals of interpolation on the latent space, for the following analyses.

For each of the ten Molearn models, the average of %_TotalErr_ decreases to 2.0 or smaller by 1000^th^ epoch (**Fig. 5B**). Meanwhile, the average of %_TotalErr_ for the MD-derived training dataset is 2.3, then being subtly larger than that for the Molearn-generated conformations. This may be due to that the training dataset is obtained under thermal fluctuation. Atomic bonds and angles should be stretched from equilibrium ones by kinetic energies at 300 K. As remarked above, the Molearn-generated conformations are energetically optimized on potential energy surface at 0 K to remove potential steric hindrance of atoms, then resulting in relatively small %_TotalErr_ values. Recalling these %_TotalErr_ values, we infer that our Molearn-generated RNA conformations have sufficient chemical accuracy. Given the chemical accuracy of the Molearn-generated conformations, we therefore consider the parameter sets at 1000^th^ epoch to be appropriate for use as representative trained models for conformation generations. Although we tested each of the ten Molearn models, we did not identify MV2003-binding-compatible conformations among those generated along the transition pathways (details are discussed in **SI-6** with **Fig. S3** and **Table S1** in **Supporting Information**). We will discuss this unexpected result in the last subsection in **Results and Discussion** in details. Then, from practical viewpoint, we examine performance of Molearn as full-atom RNA conformational generator hereafter.

We comprehensively generated TAR conformations from a set of grid points on the Molearn’s latent space (*cf.* **Fig. 2D**). The latent space of Molearn is a finite two-dimensional square region, whose x and y coordinates range from 0 to 1. A space between nearest grid points is set to 0.01 for both x and y directions. We consider 10201 (= 101 x 101) grid points on the latent space to decode into atomic structures for each Molearn model.

We confirmed the presence of MV2003-binding-compatible conformations for each set of the Molearn generated conformations. We calculated the volume of cavity formed between Cys30 and Ade35 using POVME2 ^55, 56^ (*see* **Materials and Methods**, and **SI-3** in **Supporting Information**) and selected conformations whose cavity volumes fell within the experimentally derived range (17.5-35 Å^3^; **Fig 3B**). We further screened these candidates using two interatomic distance criteria (d_2-6_ and d_4-6_; **Fig 3C**). This analysis focused exclusively on the Cys30–Ade35 region, as this region is known from NMR structures to form the MV2003 binding site; no other regions were screened for cryptic pocket formation in this study. Finally, seven of the ten independently trained Molearn models succeeded in generating TAR conformations harboring MV2003-binding-compatible cavities, while the remaining three Molearn models did not yield such conformations under the present sampling and screening criteria (**Table 1**). This outcome reflects model-to-model variability inherent to stochastic latent-space sampling rather than training failure. In total, we obtained 61 MV2003-binding-compatible TAR conformations.

**Table 1.** RMSD of Molearn-generated HIV-1 TAR conformations with potential MV2003-binding cavities relative to NMR-derived MV2003-bound TAR conformations (PDB entry: 2L8H), calculated using RNA residues C30–A35 (loop region). Unit: Å.

| Molearn model # | the number of conformation | ave. with CI95<sup>†</sup> | min. RMSd value |
| --- | --- | --- | --- |
| 1 | 8 | 3.25±0.22 | 2.78 |
| 2 | 19 | 3.04±0.11 | 2.70 |
| 3 | 2 | 2.58±0.94 | 2.10 |
| 4 | 19 | 2.77±0.06 | 2.46 |
| 5 | 1 | 2.51±0.00 | 2.51 |
| 6 | 10 | 3.06±0.43 | 2.42 |
| 7 | 2 | 2.78±0.26 | 2.64 |

<sup>†</sup>For a Molearn generated conformation, each of the NMR-derived 10 TAR conformations in the MV2003-bound state are used to calculate RMSD, then giving 10 RMSD values per one conformation.

As noted above, the training dataset does not contain any conformations with a sufficiently large cavity (*c.f.* **Fig. 4A**). Despite this limitation of the training ensemble, Molearn generates TAR conformations that extend beyond those directly sampled by MD, including structures that harbor a cryptic binding cavity. Importantly, no information from MV2003-bound TAR structures—neither cavity geometry, base-pair disruption patterns, nor ligand positions—was used during Molearn training or conformation generation; all ligand-related criteria were applied only after RNA conformations had been generated. Of the 61 conformations, we selected the structure exhibiting the minimum RMSD (**Table 1**), calculated over the loop region spanning residues C30–A35, which shows the highest structural similarity to the NMR-derived MV2003-bound TAR conformations (the relevance of this local structural similarity to MV2003 binding is discussed in the next subsection; the other conformations are shown in **Fig. S5** in **Supporting Information**). Taken together, these results demonstrate that the Molearn can access MV2003-binding-compatible TAR conformations that were not observed in previous MD-based studies ^24–26^ initiated from *apo* TAR structures, thereby providing a complementary generative capability for exploring cryptic RNA conformational states.

Nonetheless, there is a technical limitation of the present Molearn framework; its generative performance is inherently probabilistic rather than deterministic, as is typical for deep generative models. As summarized in **Table 1**, independently trained Molearn models exhibit variability in both the number of MV2003-binding-compatible TAR conformations generated and their structural similarity to the NMR-derived MV2003-bound conformations as assessed by RMSD. Importantly, all models converged during training and showed comparable loss function behavior (**Fig. 5A**), indicating that this variability does not arise from training corruption or non-convergent optimization. Instead, differences in generative performance likely reflect stochastic effects intrinsic to model initialization and data sampling (additional analyses for stochastic effects are found

in **Tables S2-6**), which also manifest as distinct latent-space organizations among trained models (**Figs. S6-9**). Achieving more stable and reproducible generation of specific functional RNA conformations therefore represents an important future challenge for extending Molearn toward a more general framework for generating previously unobserved but biologically relevant RNA conformations, such as TAR with a cryptic binding cavity.

### Functional validation of Molearn-generated TAR conformations by MV2003 docking

The 61 geometrically filtered TAR conformations were subsequently subjected to explicit docking simulations to evaluate whether they are capable of binding MV2003. This step enables us to distinguish geometrically compatible conformations from those that are functionally bindable. Hereafter, conformations whose ligand binding is explicitly validated by docking or interaction analyses are referred to as ‘bindable’. For this end, each Molearn-generated TAR conformation was docked against the NMR-derived MV2003 structures from the TAR-bound ensemble (PDB entry: 2L8H) ^21^. From each pairing of a TAR conformation with an MV2003 conformation, docking generated a set of poses (the docking procedure is illustrated in **Fig. S10**). These were ranked by the AnnapuRNA scoring function^60^, and the top-ranked pose was selected as the representative. We finally obtained 610 docked TAR–MV2003 poses. These poses were further evaluated by PoseBusters ^59^ to identify unphysical inter-atomic contacts between TAR and MV2003. Of the 610 poses, 484 satisfy the geometrical criteria and were therefore considered for subsequent analyses (*see* **SI-5** for technical details). Importantly, all 61 Molearn-generated TAR conformations yielded at least one geometrically reasonable and bindable docking pose, indicating that the excluded poses reflect unfavorable docking configurations rather than non-bindable RNA conformations.

**Fig. 6A** shows distributions of AnnapuRNA scores calculated for the 484 docked poses. The orange belt indicates the range of AnnapuRNA scores obtained from NMR-derived TAR–MV2003 complexes (−121 and −180). Approximately 86% of docking poses fall within this range, indicating that their RNA–ligand interaction patterns are comparable to those observed in experimentally resolved complexes. Furthermore, about 11% of docked poses exhibit AnnapuRNA scores lower than the minimum values observed for the NMR-derived complexes (left to the orange belt in **Fig. 6A**). These poses correspond to RNA–ligand interaction patterns that are statistically well represented in the reference dataset used to derive the AnnapuRNA score. Overall, these results indicate that TAR conformations generated by Molearn can support RNA–ligand interaction geometries comparable to those found in experimentally resolved TAR–MV2003 complexes, as assessed by the AnnapuRNA score.

**Figure 6.**
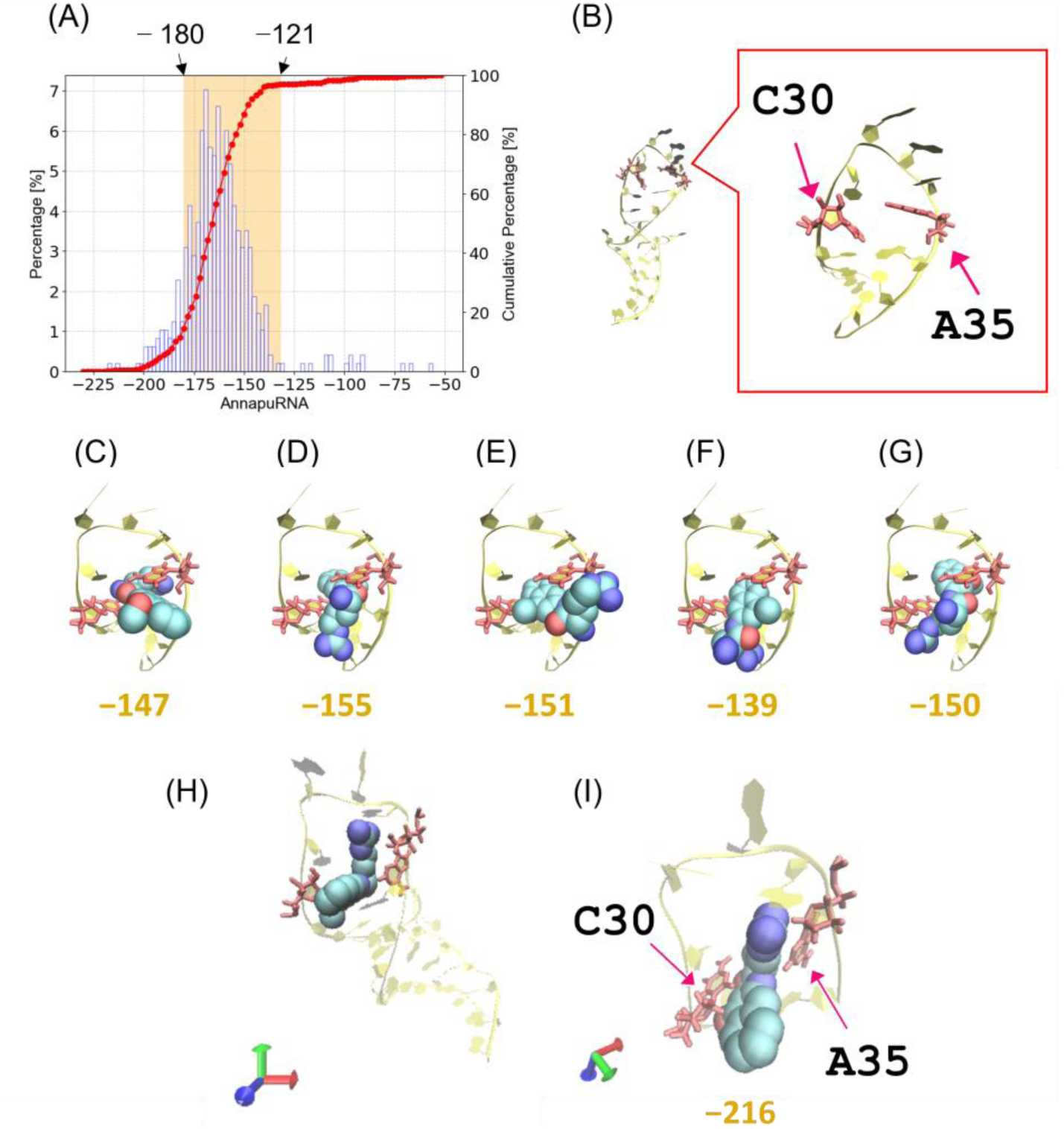
Analyses of docked HIV-1 TAR–MV2003 poses. (A) RNA-ligand binding scores calculated by AnnapuRNA scoring function for all docked poses; the orange belt indicates the value range for NMR-derived TAR–MV2003 complexes. (B) Molearn-generated TAR most similar to NMR-derived TAR in the MV2003-bound state. MV2003 interaction site is magnified in the red frame. (C)-(G) docked poses given from the conformation shown in (B). (H) Docked pose with the minimum AnnapuRNA score, obtained from other Molearn generated TAR conformation. (I) Close-up view of TAR–MV2003 conformation shown in (G). In (A), blue open bars and red closed circles are for percentages and cumulated percentage of the histogram, respectively. MV2003 and the interacting residues are shown by van der Waals spheres and red sticks, respectively. TAR structure is illustrated by transparent yellow ribbon, as in the case of **Fig. 1C**. In (C)-(G) and (I), the values shown in orange below each TAR–MV2003 docked pose denote the corresponding AnnapuRNA scores.

As remarked above, we identified one Molearn-generated MV2003-bindable conformation that most closely resembled the arrangement of the loop region spanning residues Cys30 through Ade35 observed in the NMR-derived TAR conformations in the MV2003-bound state (**Fig. 6B**). In this conformation, a cavity is present between Ade35 and Cys30 that can accommodate intercalation of the methoxy naphthalene group of MV2003, and Ade35 is positioned above Cys30 (*see* **Fig. 1C** for comparison). Docking simulations of this TAR conformation with MV2003 yielded five valid poses (the other five were excluded by PoseBusters because of steric clashes between RNA and ligand). In these poses, the methoxy naphthalene groups of MV2003 were intercalated between Cys30 and Ade35 (**Fig. 6C-G**), and the pose in **Fig. 6C** most closely resembled the NMR-derived TAR–MV2003 complex at the cryptic binding cavity (*see* **Fig. 1C**). Each of these poses received an AnnapuRNA score within the range observed for NMR-derived TAR–MV2003 complexes (−147, −155, −151, −139, and −150 in the order of panels in **Fig. 6**). Although none of these poses corresponds to the absolute lowest-scoring conformation (**Fig. 6H,** I), this likely reflects the fact that AnnapuRNA score focuses on RNA–ligand interaction patterns and is not designed to rank conformations based on global RNA structural features (*see* **SI-7** for further discussion)—they nonetheless demonstrate that Molearn-generated TAR conformations can form stable cryptic binding cavities highly consistent with the NMR-derived TAR–MV2003 complexes.

### Local functional accuracy versus global fold limitations in Molearn-generated TAR conformations

Taken together, the above results demonstrate that the present study addresses the central objective of exploring cryptic-pocket–containing RNA conformations: the generation of TAR conformations harboring a cryptic MV2003-binding cavity from *apo* conformational ensembles, without incorporating ligand-specific structural information during model training. Importantly, this case study demonstrates that Molearn can generate RNA conformations that have not been obtained by previous computational structure sampling or generation approaches, thereby addressing a long-standing challenge in RNA-targeted structure prediction within the scope of the TAR system. Recalling this milestone achieved in the present study, we next discuss the remaining challenges and limitations of the Molearn-based approach, which define the scope in which it can be most effectively applied for predicting functional RNA conformations.

Notably, we obtained TAR conformations with a cryptic binding site, forming local interactions with MV2003 as observed in the NMR-derived TAR–MV2003 complexes. It is reported that deep generative modeling-based RNA tertiary structure predictions are often less accurate in describing local interactions ^61^. Therefore, successful generation of the cryptic binding cavity of TAR highlights the technical strength of the present hybrid MD–deep generative framework implemented in Molearn, in capturing local structures (as for TAR, disruption of the base pair between Cyt30 and Ade35, shown in **Fig. 1C** and **1D**).

Meanwhile, the present version of Molearn could not consistently generate global TAR conformations that simultaneously preserve the correct overall fold while harboring the cryptic binding cavity. NMR-derived TAR conformations in complex with MV2003 adopt conformations similar to the interhelical bent conformation (*see* **Fig. S11A**; *cf.* conformation in green in **Fig. 1B**). In contrast, all 61 Molearn-generated conformations with cryptic binding cavities show substantial deviation from the interhelical bent conformation (*see* **Fig. S11B**). Although the training dataset equally includes both interhelical bent and coaxial stacking conformations (*see* **Materials and Methods**), global RNA folds of the 61 conformations do not match those of experimentally resolved MV2003-bound TAR structures (additional discussion is given in **SI-10** in **Supporting Information**). This limitation likely reflects the use of a 1D convolutional architecture operating on Cartesian coordinates, which does not explicitly preserve long-range three-dimensional spatial relationships ^62^ or rotational and translational equivariance. This observation indicates that the loss of global structural coherence is not specific to TAR, but instead reflects a general limitation of the current 1D-CNN representation when applied to extended RNA architectures. Replacing the current 1D-CNN with a geometry-aware diffusion or flow-matching model could therefore help Molearn capture long-range atomic relationships and maintain the correct global behavior of RNA structures.

### Complementary roles of Molearn relative to existing computational approaches

Molearn successfully generates TAR conformations with cryptic MV2003 binding cavity. It is then worthwhile to compare our Molearn approach with those by earlier computational studies. Studies to predict cryptic binding sites are classified into two classes, atomistic MD simulations and deep generative modeling-based RNA structure prediction, in general ^63^.

Atomistic MD simulations have been employed to search cryptic binding sites inside proteins ^64–66^, while there are few examples to straightforwardly apply them into RNA’s cryptic binding site ^26^. This discrepancy likely reflects the fact that cryptic ligand-binding sites in RNA have only recently been recognized as viable drug targets. In this context, Panei and colleagues developed SHAMAN ^26^, an MD-based approach to identify potential small-molecule binding sites in RNA by explicitly accounting for RNA conformational dynamics. By combining atomistic MD simulations with prove ligands and enhanced-sampling techniques, SHAMAN enables efficient detection of interaction-prone surface regions within sampled RNA conformational ensembles. Benchmark studies across diverse RNA systems demonstrated that SHAMAN can identify experimentally resolved binding sites, and in the case of HIV-1 TAR, it suggested a partially occluded or cryptic ligand-binding region corresponding to the MV2003 interaction site. Such MD-based detection approaches provide valuable insight into dynamically emerging pockets and local interaction hotspots within conformational space directly accessible to MD simulations, and can therefore serve as complementary tools for post-generation analyses of Molearn-generated RNA conformations.

At the same time, MD-based approaches are inherently limited to analyzing conformations sampled during the underlying MD simulations and do not aim to generate RNA conformations beyond the sampled distribution. Molearn addresses a complementary problem by generatively exploring RNA conformational space through latent-space interpolation. In the present study, Molearn was trained on *apo* TAR conformations that lacked MV2003-binding cavities, yet generated conformations harboring cryptic cavities comparable to those observed in experimentally resolved structures. Although Molearn relies on MD simulations to prepare its training dataset, the role of MD in the present framework is fundamentally different from that in MD-based cryptic-site detection approaches. In this study, MD simulations were employed only to generate locally consistent conformational ensembles around experimentally resolved equilibrium states, using relatively short simulation times, and not to directly sample cryptic pocket formation events. By generatively exploring RNA conformational space through latent-space interpolation, Molearn can access candidate ligand-bindable RNA conformations without requiring extensive MD sampling or detailed force-field optimization ^31, 67–72^, thereby complementing MD-based approaches that analyze only MD-accessible conformational ensembles.

From the perspective of inspecting cryptic binding site, it is worthwhile to note the earlier deep generative models, which are designed for proteins to predict a set of residues contributing to formation of cryptic binding sites ^73–76^. To our best knowledge, no analogous deep generative models have yet been established for RNA tertiary structure prediction. More fundamentally, cryptic binding sites in RNA molecules remains unexplored in the research field of RNA-targeted drug discovery yet. The development of RNA-specific deep-generative structure prediction model is further hindered by the limited availability of experimentally resolved RNA–ligand tertiary structures (excluding protein-RNA complexes), which constitute approximately 2.5% of all entries in PDB (cataloged by the Nucleic Acid Knowledgebase ^15^ as of January 27, 2026). Under these circumstances, the Molearn approach offers a practical route for exploring cryptic RNA binding sites by combining generative conformational sampling with external post-generation filters such as ligand-binding site detection tools (*e.g*.,SHAMAN ^26^, fpocketR ^77^)—to identify candidate binding-compatible conformations from the generated ensemble.

### Examining the scalability of Molearn using a larger RNA system

Using the paradigmatic RNA, HIV-1 TAR, as a benchmark, we demonstrated that Molearn can access RNA conformations harboring cryptic ligand-binding cavities starting from *apo* conformational ensembles. From viewpoint of technical generality, it is worthwhile to assess scalability of Molearn, here referring the ability to access cryptic, ligand-bindable conformations in larger RNAs, rather than to exhaustive recovery of bound-state geometries, we applied Molearn to a larger RNA system, the enterovirus internal ribosome entry site (IRES), which contains 41 nucleotides and is approximately 1.4-fold longer than TAR (**Fig. S13**). IRES, a well-recognized RNA drug target ^78^, binds the small molecule DMA-135 through a cryptic ligand-binding cavity that is absent in *apo* structures. Analysis of experimentally resolved IRES structures shows that DMA-135 binding is accompanied by formation of a cavity near the bulge loop region and by deviations from canonical RNA geometry, as reflected in both cavity volume and RNA-specific knowledge-based scores (**Fig. S13**). These features are not observed in available *apo* IRES structures, indicating that the DMA-135 binding site represents a cryptic pocket induced upon ligand association (**Fig. S14**).

Molearn models were trained exclusively on *apo* IRES conformations generated by atomistic MD simulations that lacked DMA-135-binding cavities (**Fig. S15**). From the generated conformational ensemble, we identified IRES conformations exhibiting cavity volumes comparable to those observed in DMA-135–bound structures. Subsequent docking simulations revealed that many of these conformations can accommodate DMA-135 with AnnapuRNA scores comparable to those calculated for experimentally resolved IRES–DMA-135 complexes (**Fig. S17**). A detailed description of Molearn model training, conformational filtering, docking procedures, and scoring analyses for IRES is provided in Section **SI-11** and **SI-12**.

Recently, Nithin *et al.* conducted a comprehensive benchmark of six RNA tertiary structure prediction methods—two deep generative models and four physics-based approaches—across 139 RNA–ligand complexes, including IRES–DMA-135 ^61^. In that study, predicted structures were evaluated based on their ability to reproduce ligand-bound geometries under uniform structural criteria. Notably, none of the tested methods recovered IRES conformations featuring the cryptic binding pocket observed experimentally (*see* **Fig. S16**). We emphasize that this comparison does not constitute a direct benchmark between Molearn and existing RNA tertiary structure prediction tools, as the objectives and evaluation protocols differ fundamentally. Whereas Nithin *et al.* focused on bound-state geometry recovery within conventional prediction frameworks, the present study explores whether a generative latent-space approach can expand *apo* RNA conformational ensembles to include cryptic ligand-bindable states. In this limited but illustrative case, Molearn accessed cryptic binding-compatible conformations for IRES that were not observed in the prior benchmark, highlighting the complementary role of hybrid MD–deep generative frameworks in RNA conformation exploration relative to existing RNA structure prediction methods.

At the same time, the present results also reveal clear limitations. Although Molearn generated IRES conformations harboring a cryptic binding cavity, these structures did not perfectly reproduce local cavity geometries and did not globally reproduce the NMR-derived IRES fold observed in the DMA-135–bound state, consistent with observations made for TAR (*see* **Tables S6** and **S7,** and **SI-12** for related discussion). A similar limitation arising from the use of a 1D convolutional architecture operating on Cartesian coordinates is observed in the larger and topologically more complex IRES system, where reduced global-fold fidelity is more pronounced. Moreover, local RNA–ligand interaction patterns in the generated IRES conformations only partially matched those of the experimentally resolved complex. These findings indicate that further methodological refinement is required for Molearn to generate RNA conformations that simultaneously capture cryptic binding sites and accurate global architectures in larger RNA systems. Directions for such refinement are discussed in the next subsection and **SI-13**.

### Technical challenges toward establishing standalone generative model for cryptic-binding site harboring RNA conformations

For Molearn to be practically applicable in RNA-targeted drug discovery, it is essential to systematically identify functionally relevant RNA conformations from the full ensemble of Molearn-generated structures. At present, however, such a detection or selection capability is not implemented within the Molearn framework itself (*see* Section **SI-6** in the Supporting Information). In particular, we observe that TAR conformations harboring the cryptic MV2003-binding cavity do not cluster near the conformational transition pathways in the latent space. They are broadly dispersed across the latent space and do not appear to occupy a distinct region separate from *apo*-like conformations, with their locations varying across independently trained Molearn models (**Fig. S6** and **S8**). This indicates that the current Molearn latent space should not be interpreted as a direct analogue of a physicochemical free-energy landscape.

Accordingly, the emergence of cryptic binding cavities in Molearn-generated conformations should be understood as a geometric extrapolation within a learned low-dimensional manifold, rather than as a direct prediction of physical transition pathways or energetic intermediates. Consistent with this interpretation, we do not observe a simple or monotonic correspondence between latent-space coordinates and geometric descriptors of cavity formation, such as cavity volume or inter-base distances. This reflects the fact that Molearn’s latent variables are optimized for structural reconstruction and smooth interpolation, rather than for direct physical or functional interpretability. Developing latent representations that can be directly aligned with functional metrics remains an important direction for future methodological refinement.

In this sense, Molearn does not hallucinate arbitrary structures, but instead recombines correlated structural features present in the MD-derived training ensemble in a continuous and self-consistent manner. Nonetheless, developing an intrinsic and prior-free strategy to identify functional RNA conformations within the Molearn latent space remains an important direction for future refinement. Achieving this would not only improve practical applicability, but could also provide deeper insight into the key descriptors underlying RNA functional conformational changes.

Considering success of advantageous framework in the field of deep generative modeling-based biomolecule’s conformation prediction ^79, 80^, incorporating geometry-aware architectures—such as diffusion-based models with E(3)-equivariant representations—into Molearn while retaining its central concept of transition-path generation could substantially improve generative performance. Such extensions may enhance the accuracy of both local molecular features, including cryptic ligand-binding sites, and global RNA architectures, thereby broadening the applicability of Molearn-based approaches to RNA-targeted structure-based drug discovery.

## Concluding remarks

Cryptic binding sites in RNA represent promising drug targets because of high specificity and low off-target potential. However, RNA-targeted structure-based drug design (SBDD) remains technically challenging, owing to the limited availability of experimentally resolved RNA tertiary structures and the inherent difficulty of predicting cryptic binding sites from *apo* conformations. In this study, we addressed this challenge by extending the applicability of the hybrid MD–deep generative framework, Molearn to predict RNA conformations harboring cryptic binding sites. Starting exclusively from *apo* RNA conformations, Molearn generated RNA conformational ensembles that included structures containing cryptic ligand-binding cavities in both the HIV-1 TAR and IRES systems. In contrast to prior MD- or deep generative modeling-based studies that primarily analyze conformations directly accessible within sampled ensembles, the present work demonstrates that a hybrid MD–generative strategy can expand *apo* RNA conformational space to access ligand-binding-compatible cryptic states that were not observed in corresponding *apo* MD trajectories. This capability provides a conceptual and methodological baseline for future developments in RNA generative modeling. At the same time, key challenges remain in routinely generating RNA conformations that simultaneously recover cryptic binding sites and accurate global folds across RNAs of varying length and complexity. Addressing these challenges will require further refinement of the Molearn framework and related generative models, with the ultimate goal of establishing robust deep generative approaches for RNA-targeted structure-based drug design (SBDD).

## Author contributions

M.H. supervised the project. I.K. conceived and designed the research, performed computational experiments and analyzed all data. I.K. and M.H. interpreted and discussed the results, wrote the manuscript, and reviewed the manuscript.

## Conflict of interest

The authors declare no competing interests.

## Data availability

The MD simulation dataset, Molearn-related data and cavity analyses for predicted RNA structures in the earlier study^61^ that support the findings of this study are obtained from and https://waseda.app.box.com/folder/300778804035?v=data-molearnA. All codes needed to run RNA-adjusted Molearn are obtained from https://github.com/hmdlab/molearnA.

The supplementary information is available free of charge. Further details for atomistic MD simulations, additional physicochemical analyses for NMR-derived and Molearn-generated conformations, supporting Tables and Figures.

## Acknowledgements

This research was supported by the Core Research for Evolutional Science and Technology (CREST) under Grant Number JPMJCR21F1, Japan Agency for Medical Research and Development (AMED) under Grant Number 23ae0121049h0003, and Grant-in-Aid for Scientific Research (A) under Grant Number 23H00509. The computation associated with Molearn was performed using Research Center for Computational Science, Okazaki, Japan (Project: 23-IMS-C129; 24-IMS-C123; 25-IMS-C122). ChatGPT (OpenAI) was used exclusively for language editing and clarification of sentence expression during manuscript preparation. After using this tool, the authors reviewed and edited the content as needed and take full responsibility for the content of the publication.

## Notes

### Competing Interest Statement

The authors have declared no competing interest.

### Summary of Updates

Modify the title and terminology in the main text.

