## Supporting_Information for "Hybrid molecular dynamics–deep generative framework expands *apo* RNA ensembles toward cryptic ligand-binding conformations: application to HIV-1 TAR"

[*]Ikuo Kurisaki:

[*]Michiaki Hamada:

#### SI-1. Structural characterization for other two sets of *apo* TAR conformations.

The three sets of atomic coordinates of *apo* TAR are registered in Protein Data Bank (PDB entries: 7JU1, 8THV and 1ANR). In this study, we chose the 7JU1 because it exhibits lower BRiQ scores, indicating more canonical *apo* TAR conformations, compared with 1ANR or 8THV. 1ANR ensemble shows higher BRiQ scores than the conformations in the MV2003-bound state (PDB entry: 2L8H) (**Fig. 3A** and **S1A**). The 8THV ensemble exhibit BRiQ scores similar to those of 7JU1; however, the model No. 19 in 8THV shows deviation from canonical RNA geometry comparable to those observed in the MV2003-bound conformations.

1ANR exhibits greater BRiQ scores than 7JU1, 8THV and 2L8H, maybe reminding us of presence of MV2003-binding compatible cavity. However, any of the conformations does not have MV2003-binding cavity. Two of twenty conformations of 1ANR, No. 5 and No. 14, show expected volume values, but they do not satisfy the distance-pair condition (*see* **Fig. S1B** and **S1C**). As shown in the left and center panels, each of their Ade35 is flipped out from the loop. No. 17 satisfies the distance pair condition (**Fig. S1C** and the right panel in **Fig. S1D**) but does not form an MV2003-binding compatible cavity. Accordingly, it can be said that there are no NMR-resolved *apo* TAR conformations with the MV2003-binding compatible cavity.

#### SI-2. Details of MD simulation procedure

After MM simulations, the system temperature and density were relaxed through the following five MD simulations: NVT (0.001 to 1 K, 0.1 ps) → NVT (1 K, 0.1 ps) → NVT (1 to 300 K, 20 ps) → NVT (300 K, 20 ps) → NPT (300 K, 300 ps, 1 bar). In each of MM and MD simulations, the atomic coordinates of non-hydrogen atoms in TAR were restrained by the harmonic potential with force constant of 10 kcal/mol/Å$^2$ around the initial atomic coordinates. The first two NVT MD simulations and the other MD simulations were performed using 0.01 fs and 2 fs for the time step of integration, respectively. The first and second NVT MD simulations were performed using Berendsen thermostat[3] with a 0.001 ps of coupling constant. Meanwhile the following three simulations were performed using Langevin thermostat with 1-ps$^{-1}$ of collision coefficient. In the first NVT MD simulation, the reference temperature was linearly increased along the time-course. In the NPT MD simulation, the system pressure was regulated with Monte Carlo barostat, where the system volume change was attempted by every 100 steps. Each set of initial atomic velocities was randomly assigned from the Maxwellian distribution at 0.001 K.

For each atomic coordinates obtained above, an RNA conformation also was structurally relaxed in aqueous solution through the following 7-step MD simulations: NVT (0.001 to 1 K, 0.1 ps) → NVT (1 to 300 K, 0.1 ps) → NVT (300 K, 10 ps, 10 kcal/mol/Å) → NVT (300 K, 40 ps) → NVT (300 K, 40 ps) → NVT (300 K, 40 ps) → NPT (300 K, 1 bar, 200 ns). The first two NVT MD simulations and the other MD simulations were performed using 0.01 fs and 2 fs for the time step of integration, respectively. In the first two NVT simulations, the reference temperature was linearly increased along the time-course. In each NVT simulation, temperature was regulated using Langevin thermostat with 1-ps$^{-1}$ collision coefficient. In the last 200-ns NPT simulation, temperature and pressure were regulated by Berendsen thermostat [1] with a 5-ps coupling constant, and Monte Carlo barostat, where system volume change was attempted by every 100 steps, respectively. The initial atomic velocities were randomly assigned from the Maxwellian distribution at 0.001 K.

Employing the MD simulation protocol described above, we obtained three pairs of 200-ns MD trajectories initiated from interhelical bent and coaxial stacking conformations (**Fig. S2A–B**, **Fig. S2C–D**, and **Fig. S2E–F**). In all trajectories initiated from the interhelical bent conformation, stable RMSD profiles were observed throughout the simulation time (**Fig. S2A**, **S2C**, and **S2E**). For trajectories initiated from the coaxial stacking conformation, two trajectories exhibited apparent changes in RMSD relative to the initial structures around 120 ns (**Fig. S2B** and **S2D**), indicating transitions away from the starting conformations. The remaining trajectory retained stable RMSD profiles relative to both interhelical bent and coaxial stacking reference structures over the 200-ns time window (**Fig. S2F**), although similar transitions may occur on longer timescales. These observations are consistent with intrinsic conformational flexibility of TAR in the coaxial stacking state under thermal fluctuations. For Molearn model training, we selected the pair of MD trajectories shown in **Fig. S2A** and **S2B**, which together capture both stable interhelical bent conformations and conformational transitions originating from the coaxial stacking state, while maintaining overall structural integrity. The additional MD trajectories shown in **Fig. S2C–F** were generated to assess the robustness and reproducibility of the RMSD behavior under the same simulation protocol and were not used for Molearn model training.

#### SI-3. Cavity volume calculations with POVME2

Volume of MV2003-binding cavity was calculated by using POVME2 [2,3] with the following conditions. The inclusion region is a sphere whose radius and center are 3 Å and the midpoint between $C_1$' atom of Cyt30 and $C_1$' atom of Ade35. The exclusion region is a sphere whose radius and center are 4 Å and the midpoint between the phosphate atoms of Ura31 and Gua34. The latter region works to filter out TAR conformations without proper rearrangement of Cyt30 and Ade35 (an example is shown in **Fig. S1D**, which form non-Watson Click hydrogen bonding between Cyt30 and Ade35, and has a cavity over the paired bases). Grid space and distance cutoff from atoms in RNA is set to 0.5 Å and 1.09 Å. Isolated points which do not have three or more neighboring points are removed from a set of volume element.

#### SI-4. RNA−ligand docking simulation with RLDOCK

We performed TAR-MV2003 docking simulations with RLDOCK program suite [4]. The 61 Molearn-generated MV2003-binding-compatible conformations were employed (*c.f.* **Table 1**). We obtained ten sets of atomic coordinates of MV2003 from the NMR-resolved conformations (PDB entry: 2L8H) and used each of them for the RNA−ligand docking simulations. Aiming to test whether MV2003 can access a cavity, we consider local docking simulation with a given docking site. A docking site is a cube whose size and center are 20 Å and the midpoint between $C_1$’ atom of Cyt30 and $C_1$’ atom of Ade35, respectively. Atomic charges for RNA and ligands are assigned following the original study of RLDOCK[4], using AMBER14SB force field [5] for RNA and Gasteiger charges for ligands [6]. Atomic coordinate files are arranged by Open Babel (version 3.1.1) [7] to execute docking simulation with RLDOCK. Generated docked poses were evaluated using AnnapuRNA scoring function [8] and the top ranked structure is analyzed as a representative TAR–MV2003 docking pose.

#### SI-5. Evaluation of docked TAR–MV2003 poses

Interatomic steric clashes between ligand and Molearn-generated RNA in docking poses were evaluated by using the PoseBusters program [9]. A geometrical relationship of atom pair is classified with two categories, severe and acceptable. Calculating an inter-atomic distance, $d$, we compared $d$ with sum of van der Waals radii of atoms, $s$. If $d < 0.7$ [Å], we consider that atoms are severely clashed. When any severe steric clash is found, the corresponding docked TAR–MV2003 pose is ignored in the following analyses. The value of 0.7 [Å] is derived from the geometrical criteria discussed in the original study developing PoseBusters, that is, *internal steric clash*.

#### SI-6 Challenge to systematically detect TAR conformations with the cryptic binding cavity

In this study, Molearn generated a diverse ensemble of TAR conformations, from which candidate MV2003-binding-compatible structures were identified using post-generation geometric criteria. However, the systematic identification of functionally relevant conformations directly from the Molearn-generated ensemble—without applying ligand-specific geometric filters—remains an open challenge.

Supposing that ligand-bindable RNA conformations are deviated from canonical RNA geometry by disrupting intra-RNA interactions, we expect to find RNA conformations of interest on conformational transition pathways connecting basins in an *apo* RNA ensemble. This is the working hypothesis and motivation to use Molearn. As discussed in **Introduction**, it is computationally expensive and sometimes challenging task for atomistic MD simulations to examine a transition pathway. To overcome this inevitable technical limitation of MD simulations, we expected that Molearn enable us to test the above hypothesis by analyzing Molearn-derived conformations along pathways.

The hypothesis has one important assumption that we explore plausible RNA conformation transition pathways from Molearn's latent spaces. By using the Molearn models at 1000$^{th}$ epoch, a set of TAR conformations were generated on linearly interpolated 48 points connecting two the representative conformations (similar to No. 4 and No. 20 remarked in **Fig. 1B**, a pair of MD-derived conformation with the maximum RMSD value). Following the original Molearn study but recalling the complexity of all-atom RNA structure, we used 48 points as test. After energetically refining the generated conformations with the RNA-featured molecular mechanics simulation program QRNAS [10], we calculated BRiQ-score [11] profiles along the pathways (**Fig. S3A-J**).

Each of the ten models often generates conformations with BRiQ scores over 50 [$k_BT$]. It is noted that, for the NMR-derived MV2003-bound conformations, the maximum BRiQ score is about 50 [$k_BT$] (*see* **Fig. 3A**). Molearn-generated conformations appear to be relatively deviated from canonical RNA geometry compared with the NMR-derived ones. Besides, we found remarkably large value of BRiQ values on pathways, *e.g.,* 321 at the 15$^{th}$ frame in **Fig. S3A** (*see* **Fig. 3A** for comparison of BRiQ score between *apo* and MV2003-bound states). Such large BRiQ peaks suggest the presence of TAR conformation fairly deviated from canonical RNA geometry. These results may not be simply due to imbalance between BRiQ scores for pathway analysis and AMBER-based potential energy function [10] for structure refinement. Nonetheless, since a BRiQ score at the peak is distinguishably large compared with others, the pathways are not minimum BRiQ score-based pathways along which TAR undergoes conformational change. This disappointed result of building transition pathways in Molearn latent space infers that the present Molearn model is still immature to reproduce plausible transition pathways of biomacromolecule for the *all-atom* level resolution model.

Nonetheless, from a practical viewpoint, it is still worthwhile to test the performance to generate TAR conformations harboring the cryptic binding pocket. In the pathways shown in panels **A**, **D**, **E** and **F** in **Fig. S3**, we found seven TAR conformations (annotated by the numbers from **1** to **7** in **Fig. S3**) satisfying the cavity volume criterion. Further, the two of the seven satisfies the other criterion for base-pair distance, too (**Fig. S3K**). One of the two conformations annotated by **2** and illustrated in **Fig. S3L** has a cavity between Cyt30 and Ade35 (**Fig. S3K**) but appears to be far from the NMR-resolved one; the positional relation between Cyt30 and Ade35 is different from that observed in the NMR-derived MV2003-bound conformations (compared with that shown in **Fig. 1C**). The remaining one annotated by **6** still suffers from steric hindrance around the loop region (**Fig. S4**), thus being chemically improper for subsequent atomic-level analyses. Then, we concluded that these two conformations are not promising MV2003-binding-compatible ones and did not use them for further analyses (*i.e.*, RNA−ligand docking simulations and RNA−ligand interaction pattern analysis) for the two conformations. Accordingly, we could not accommodate to systematically generate MV2003-binding-compatible TAR conformations by interpolation between the representative TAR conformations.

This disappointed result might come from a technical process to TAR generate conformations, a definition of transition pathways. We linearly interpolated the two structures on the latent space as in the original Molearn study, but it is not sure that such a transition pathway run on an optimal route, composed of canonical RNA structures.

Notably, the original study only considered further simple molecular models which have the four backbone atoms and one sidechain atom ($C_\alpha$, C, N, O, $C_\beta$) for any amino acid residue, and global domain motions are examined. Meanwhile, we use full-atom RNA structures (27 heavy atoms per RNA residue, comprising phosphate, sugar and base) to train Molearn models. TAR undergoes complicated, multi-scale conformational changes (*cf.* **Fig. 1B**), through which TAR takes both global and local conformational rearrangements. Full-atom description of RNA is inevitable for subsequent application to SBDD, while the shapes of knowledge-based score profiles are substantially jagged and volatile. Besides, we found spiky changes for each of BRiQ score-based profiles and relatively large scores at the points (quantitatively characterized in **Table S1** in Supporting Information). It is noted that biomacromolecules inherently have rough BRiQ score profiles on their conformational transition pathways [12-14]. The RNA model employed in this study may lead to rather complicated profiles along transition pathways, compared with the simplified one employed in the original Molearn study [15]. Recalling this observation, we concluded that the linearly interpolated points are away from plausible RNA conformation transition pathways.

Then, we constructed a minimum BRiQ score pathway by using Dijkstra's algorism (orange trace in each panel in **Fig. S6**). As is the above case of the linear interpolation, we use the representative interhelical bent and coaxial stacking conformations as each of the two end points. A minimum score pathway is assigned to grid points in the Molearn latent space (*see* **Fig. 1D**). Any of pathways is found inside the convex polygon (dotted line in **Fig. S6**). This may be explained by considering the formula of loss function (**Eq. 1**), which has both reproduction of the input dataset and physicochemical accuracy of interpolated conformations (*see* **Fig. 2C**). Indeed, conformations inside the convex polygon have relatively small values of $\%_{\mathrm{TotalErr}}$ and exhibit relative similarity to canonical RNA geometry in terms of BRiQ score (*see* **Tables S2** and **S3**). Compared with the interpolation-generation pathways, both maximum value and spikiness of BRiQ score-based profile are apparently decreased in these pathways (*see* **Fig. S7**, **Table S1** and **SI-8** for related discussion). Therefore, using these pathways may be better to test our conjecture than using the above linearly interpolated pathways.

We then tested whether MV2003-bindable conformations (red squares in **Fig. S6**) appear through conformational transition between the two representative interhelical bent and coaxial stacking conformations. Despite our expectation, they are not necessarily found near these minimum score pathways. As for the models shown in **Fig. S6B, S6D** and **S6F**, a part of conformations seems to match our expectation, although the relative positions of these conformations to the corresponding minimum score pathways are different among the three models. Thus, appearance of MV2003-bindable conformations near the minimum score pathways is accidental and our Molearn latent spaces may be impractical to systematically discover such conformations based on our conjecture, so far. This situation was unchanged even if we used a different pair of starting and ending points to make transition pathways (*see* **Fig S8** and **S9** and the related discussion in **SI-9** in Supporting Information). Latent spaces of Molearn models could not inform where ligand-bindable conformations appear yet.

In summary, it may be still unfeasible to systematically discover such conformations by analyzing a BRiQ score-based landscape of *apo* TAR on the latent space. To detect RNA conformation of interest from the latent space, we need to improve Molearn framework. It is pursued but elusive to construct physically-reasonable free energy landscapes of interest by using biomolecular structure-generative models [16] and improvement of Molearn's performance is categorized as this kind of problem. We consider such works as the next steps of this study.

#### SI-7 Structural variation of docked TAR–MV2003 poses and remarks for evaluation of docked-pose ranking

Using Molearn-generated TAR conformations, we obtained a diverse set of docked TAR–MV2003 poses. In **Fig. 6C**, the orientation of MV2003 within the TAR cryptic binding cavity is similar to the experimentally observed one (*see* **Fig. 1C**). Meanwhile the other four appears to adopt opposite orientations except for the pose shown in **Fig. 6C**. This could be attributed to RNA−ligand docking simulations. It is not the main subject of this study to investigate docking mechanisms through a comprehensive comparison of docking simulation protocols for TAR and MV2003, so that we did not consider this issue further.

None of the five docked poses shown in **Fig. 6C-G** is the docked TAR–MV2003 pose with the minimum AnnapuRNA score (−216), which shown in **Fig. 6H** and **I**). In this lowest-scoring conformation, the methoxy naphthalene group of MV2003 is intercalated between Cyt30 and Ade35, but the relative position between Ade35 and Cys30 is apparently different from that of the NMR-resolved TAR in the MV2003-bound state (**Fig. 6I**; *see* also **Fig. 1C** for comparison). This pose could be identified as the lowest-scoring conformation because the docked poses are evaluated solely based on local RNA–ligand interaction pattern only. Indeed, it is thus elusive whether the docked TAR–MV2003 pose with the lowest-AnnapuRNA score is more plausible than the above five docked poses.

We here employed AnnapuRNA as a representative scoring tool because there are no available RNA-specific score functions which comprehensively consider RNA, ligand

and RNA–ligand interaction patterns so far. Although it is technically possible to calculate free energy by using MD simulations, this approach requires substantial computational cost and careful selection of empirical force fields. However, further evaluation of these docked poses is beyond the scope of this study, so that we do not discuss this point further. This issue will be examined once a suitable scoring function to reliably estimate RNA–ligand binding free energies becomes available. Accordingly, AnnapuRNA scores should be interpreted as indicators of local RNA–ligand interaction plausibility for docked poses, rather than as metrics for assessing global RNA structural fidelity or overall functional relevance.

#### SI-8. Evaluation of spikiness in BRiQ score profile

A BRiQ score profile calculated for a given pathway often has a spiky point (**Fig. S3** and **Fig. S7**), which corresponds to remarkable deviation from canonical RNA geometry through conformation transitions. A pathway with smaller spikiness would be more employed for RNA conformational transitions. Recalling the above observations, we quantify spikiness in a profile with the following formula.

$$max(\{E_i\}_P) - average(\{E_i\}_P)$$

$P$ at the right of $\{E_i\}$ means that values are given from points on a transition pathway. The subscript $i$ and $E_i$ denote index for a TAR conformation on the transition pathway $P$ and a BRiQ score for the $i^{\text{th}}$ conformation, respectively. In summary, $\{E_i\}_P$ denotes a set of $E_i$ considered for $P$. This metric becomes relatively large when a spikiness is apparent.

#### SI-9. Optimal pathway with alternative TAR conformation pair

Aiming to construct better BRiQ score-based profiles, we used an alternative pair of

TAR conformations to search for a minimum BRiQ score pathway. Relatively low-scored conformations correspond to basins in the profile and are expected to appear as denser regions in conformational coordinate space. Two subsets of MD-derived conformations were projected onto the Molearn latent space, and representative conformations were selected from relatively dense regions. One subset was obtained from the MD simulation initiated from model No. 4 of 7JU1. This subset was clustered on the latent space using the Density-Based Spatial Clustering of Applications with Noise (DBSCAN) algorithm, with distance threshold 0.05 and minimum sample number 5 as test parameters. For the densest cluster, the conformation closest to the cluster centroid was identified as one terminal conformation. The other subset, derived from the MD simulation starting from model No. 20 of 7JU1, was processed in the same manner to obtain the other terminal conformation. Using this alternative pair, we calculated a minimum-BRiQ score pathway (**Figs. S8** and **S9**); however, MV2003-bindable conformations were not identified along this pathway.

#### SI-10. Evaluation of global conformational plausibility of Molearn-generated TAR structures

We evaluated the global structural plausibility of Molearn-generated TAR conformations using the BRiQ score, a statistical potential that reflects canonical RNA tertiary geometry. Using the NMR-resolved TAR conformations (*see* **Fig. 3A**) as a baseline of the score, we first analyzed the 400 MD-derived snapshot structures employed as the Molearn training dataset (the leftmost violin plot in **Fig. S12**). The BRiQ score distribution of these MD snapshots is broadly comparable to those calculated for the NMR-resolved *apo* and MV2003-bound TAR conformations (*see* **Fig. 3A** and **Table S4**), although it is slightly shifted toward higher values relative to the NMR-resolved *apo* ensemble. This shift likely arises because the MD snapshots sample thermally fluctuating conformations under an empirical molecular mechanics potential and therefore deviate modestly from the initial NMR structures.

In contrast, TAR conformations generated by Molearn models over the entire latent-space grid exhibit substantially broader BRiQ score distributions, with markedly larger average and maximum values (the ten central violin plots in **Fig. S12**). This behavior reflects the original design of Molearn, which was developed to explore transition pathways between two biomolecular states. The Molearn loss function includes a *path term* (**Eq. 1**) that incorporates a physicochemical energy function and enforces local structural validity, such as reasonable bond lengths and angle bending, along the interpolated pathway. However, this term does not explicitly encode long-range tertiary constraints required to reproduce native RNA global conformations. Consequently, satisfying the loss function does not guarantee accurate reconstruction of global RNA folds, particularly for structures sampled off the learned pathway, even after subsequent energetic refinement using molecular mechanics simulations.

When the analysis is restricted to the 61 MV2003-bindable conformations (the rightmost violin plot in **Fig. S12**), the BRiQ score distribution becomes substantially narrower, indicating that these conformations do not suffer from severe global structural distortions. Nevertheless, their BRiQ scores remain higher than those of the NMR-resolved MV2003-bound TAR conformations. This result suggests that, while Molearn-generated MV2003-bindable conformations achieve locally appropriate geometries compatible with ligand binding, they do not yet fully reproduce the native-like global tertiary organization of the experimentally resolved ligand-bound TAR structures.

#### SI-11. System setup to Molearn training and structure analyses for *apo* IRES

We obtained the two sets of 10 NMR-resolved *apo* enterovirus Internal Ribosomal Entry Site (IRES) structures (Protein data bank (PDB) entries: 5V16 and 5V17) as the conformation ensemble in the ground state [17,18]. We considered both 5V16 and 5V17 because they are different in global conformations, resembling coaxial stacking and interhelical bent conformations of TAR. The 5V17 is a mutant construct so that each of conformations are reverse-mutated to match the wild type RNA sequence and energetically optimized on the reverse-mutated residues in vacuum. Among each of the 10 conformations, the models No. 10 of 5V16 and No. 6 of 5V17 are selected as the representative coaxial stacking and interhelical bent conformations, respectively (*see* **Fig. S13B**) because they are the most structurally different pair among the 190 pairs (*i.e.*, ${}_{20}C_2$) in terms of RMSD.

Each of the two *apo* IRES structures are solvated in the rectangular box with 22300 water molecules, and electronically neutralized with 40 $Na^+$. Force field parameters are assigned with each of atoms as in the case of TAR and 500-ns MD simulations are performed under similar computational conditions (*see* **Materials and Methods** in the main text), where snapshot structures are recorded by 1 ns. As for each of 500 snapshot structures, we picked up ones with incompetent ligand (DMA-135)-binding cavity (the volume values are found within the range of those of *apo* IRES conformations, 0 to 188 Å$^3$; *see* **SI-11** for detailed analyses). Finally, we obtained 395 and 392 conformations for 5V16 and the reverse-mutated 5V17, respectively.

The training dataset is made from the 787 snapshot structures by removing water and ion atoms. Each snapshot structure is superposed on the models No. 10 of wild type IERS (PDB entry: 5V16) by using root-mean-square fitting algorithm with phosphate and C4' atoms in the stem region (residues number are from 128 to 131 and from 165 to 168 in biological annotation, *see* **Fig S13A**). We performed ten independent trainings of Molearn model with this dataset, with the similar training procedure (*see* **Materials and Methods** in the main text).

Volume of DMA-135-binding cavity was calculated by using POVME2 [2,3] with the following conditions. The inclusion region is a sphere whose radius and center are 7 Å and the center of gravity among $C_1$' atoms of Ade133, Ade136 and Ade139, respectively. The other computation conditions are similar to those for TAR.

We performed IRES–DMA-135 docking simulations using the RLDOCK program suite [4]. All Molearn-generated DMA-135-binding-compatible conformations were employed (*see* **Table S5**). Ten DMA-135 conformations were extracted from the NMR-resolved IRES–DMA-135 complexes (PDB entry: 6XB7) and each was docked separately to the Molearn-generated IRES conformations. Aiming to test whether a DMA-135 can access a cryptic binding cavity of IRES, we consider local docking simulation with a given docking box, whose size and center are 10 Å and the center of gravity among $C_1$' atoms of Ade133, Ade136 and Ade165, respectively. The other computation conditions are similar to those for TAR. Generated docking poses were evaluated using AnnapuRNA scoring function [8] and the top ranked structure is analyzed as a representative IRES–DMA-135 docking pose.

#### SI-12. Generation of cryptic binding site in IRES

We considered scalability of Molearn to generate functional RNA conformations by using alternative RNA structure, IRES. This RNA consists of 41 RNA residues (**Fig. S13A**), thus being 1.4-folod longer than TAR in terms of sequence length [17]. IRES takes two distinctive conformations [17,18] (**Fig. S13B**) as well as TAR, and the DMA-135-binding site is similarly *cryptic* (**Fig. S13C**; explained in detail below). Meanwhile, IRES is different from TAR in the biological function and ligand-binding site. TAR regulates RNA transcription while IRES accelerates translation initiation of mRNA. The ligand binding site of IRES is found around bulge loop region (that in TAR is found in the stem loop, *see* **Fig. 1C**). Thus, considering IRES is worthwhile to test extensive applicability of Molearn to generation of RNA conformations with cryptic binding sites.

There are the three available PDB entries for IRES, which are wild type *apo* form, mutated *apo* form and wild type DMA-135-bound form (5V16, 5V17 [17] and 6XB7 [18], respectively). None of wild type and mutate *apo* forms has DMA-135-binding pockets as elucidated below. Aiming to globally sample conformations with Molearn, we reverse-mutated each model of 5V17 (annotated by Rev. Mut, hereafter) and employed the following analysis, with expecting to expand repertoire of *intermediates* conformations generated by Molearn.

IRES binds to DMA-135 with a cryptic binding pocket. We show it by analyzing two structural descriptors, DMA-135-binding cavity volume and RNA-specific knowledge-based score, for *apo* conformations and those in the DMA-135-bound state (**Fig. S13D** and **S13E**, respectively). DMA-135 binding accompanies expansion of intra-IRES cavity around the bulge loop (**Fig. S13D**). Comparing BRiQ scores of 5V16 with those of 6XB7, we find that the DMA-135-binding deviates IRES structures from canonical RNA geometry (**Fig. S13E**) as in the case of TAR. This observation is consistent with our expectation, remarked in **Introduction** in the main text, that RNA binds to ligands in by disrupting intra-RNA interactions.

The volume of DMA-135-binding-compatible cavity basically is larger in liganded forms than in *apo* forms (**Fig. S13D**). Nonetheless, one liganded-IRES conformations show smaller volume values than *apo* ones. The No. 6 of 6XB7, IRES in the DMA-135-bound state, has relatively small cavity volume, compared with the No. 6 and 7 of 5V17 (Rev. Mut.). This is due to that IRES can take alternative DMA-135-binding mode without DMA-135 access into the cavity, directly bound on the surface of IRES (**Fig. S13F**). Except for the surface-contact DMA-135-binding modes, which are found in models No. 2, 6, 7 and 8 of 6XB7, the DMA-135 binding cavity takes lager volume in DMA-135-bound states than in *apo* states. Since we are interested in formation of cryptic binding sites and we do not consider such a surface-contact DMA-135-binding model. Besides, the four RNA residues, Ade133, 136, 139 and 142 undergo a rearrangement upon binding to DMA-135 (*see* **Fig S14**). Then, we infer that the IRES's DMA-135-binding cavity is a cryptic site induced by DMA-135-binding and the volume range of cryptic binding cavity from 239 to 373 Å$^3$ in DMA-135-bound states (*see* the orange belt in **Fig. S13D**).

Next, we examined BRiQ scores for the experimentally-resolved IRES conformations. Model No. 8 of 5V17 (Rev. Mut.) shows relatively large scores compared with several models of 6XB7 (DMA-135-bound). This may be due to use of mutation construct to solve the RNA structures and our MM simulations have not sufficiently relaxed the conformation toward wild type *apo* forms. Nonetheless, this point is irrelevant to the following discussion, because we do not use Model No. 8 of 5V17 for the following conformational sampling because of alternative DMA-135 binding mode (*see* **Fig. S13F**). Upon sampling IRES conformations with MD simulations, we consider No. 10 of 5V16 and No. 6 of 5V17 (Rev. Mut.) owing to the largest RMSD value among the possible pairs of *apo* conformations. As in the case of TAR, we consider such a pair to prepare two distinct conformation ensembles for IRES.

We performed a 500-ns NPT MD simulation for each of the two *apo* IRES structures (*see* **SI-10** for details of simulation procedure) and obtained two set of 500 snapshot structures. Differing from the case of TAR, a part of IRES conformations shows cavity volume larger than the upper bound for *apo* IRES, 188 Å$^3$. Then, we excluded such conformations and finally obtained 787 conformations for Molearn's training dataset (395 and 392 conformations come from MD simulations for 5V16 and reverse mutated 5V17, respectively).

Using this training dataset and a similar training protocol used for TAR, Molearn models for IRES were trained 10 times, independently. Molearn models were trained by using 5 nodes on CPU machines (AMD E7763) of RCCS and took around 8.6 days for a single 1500 epoch training. The total number of trainable network parameters is 5414899, which occupies disk space of 581 MB (the network parameter sizes are the same between TAR and IRES, due to use of the same Molearn architecture, *see* Materials and Methods; the larger disk space usage comes from greater size of IRES than that of TAR, which influences memory size for loss function terms *etc*.). A network loss stably decreases to seemingly reach convergence by training 1500 epoch for each Molearn model (**Fig. S15**). As remarked in the main text, the present version of Molearn works as a *simple* conformation ensemble generator. Then, we straightforwardly use each of the trained Molearn models at 1500 epoch to generate IRES conformations from grid points on the latent space.

As in the case of TAR, we generated 10201 IRES conformations for each of models.

They were energetically optimized by using QRNAS with a similar procedure for TAR (*see* Materials and Methods). A QRNAS computation for IRES takes about 85 minutes with single core of the RCCS's CPU machines and completing all 10121 QRNAS computations needs 29 hours with 500 parallel computations in RCCS. Among the QRNAS-optimized conformations, we picked up those with $\%_{\mathrm{TotalErr}}$ of less than 2.3, as well as TAR, to ensure chemical plausibility of IRES structure. Then, among the 102010 Molearn-generated IRES conformations, derived from the 10 trained models in total, 84842 conformations are filtered. We analyzed cavity volumes for these filtered conformations and detected those whose volume values are found within the value range of IRES in the DMA-135-bound state, from 239 to 373 Å$^3$. From cavity volume filtration, we obtained 3938 conformations for RNA−ligand docking simulations.

For each of 3938 conformations, we test the 10 independent DMA-135 conformations, obtained from the NMR-derived IRES–DMA-135 complexes (39380 docking simulations were executed to make sets of IRES–DMA-135 docked poses). Each set of docked poses is ranked by AnnapuRNA scoring function and one with the lowest score is selected as the representative. The lowest-scored poses are, further, evaluated using PoseBusers for chemical plausibility of RNA−ligand interaction patterns. We finally obtained 12419 poses via the filtration. In terms of AnnapuRNA scores, 95% of IRES–DMA-135 docked poses are found within the value range calculated for NMR-resolved IRES–DMA-135 complexes (**Fig. S17A**). The remaining 5% shows lower AnnapuRNA scores. Taken together, these results indicate that a large fraction of Molearn-generated IRES conformations that pass the geometrical filtering criteria can form docked complexes with DMA-135 exhibiting RNA–ligand interaction patterns comparable to experimentally resolved structures, thereby validating their ligand bindability at the docking level.

As for IRES, Nithin and colleagues addressed to reproduce conformations in DMA-135 bound state but did not accomplish those harboring the cryptic binding cavity. They tested ability of six RNA structure prediction methods to recover RNA conformations in ligand-bound states, where IRES (PDB entry: 6XB7) is found as one of the target 139 RNA–ligand complexes. Then, we examined whether these conformations have DMA-135-binding-compatible cavity. Notably, none of these structures has a cavity with ligand-binding-compatible volume (**Figure S14A**). The models generated by RNA BRiQ [11] and Rhofold [19] show 'cavity volumes' larger than 373 $Å^3$, upper bound of cavity volume obtained from the NMR-derived IRES structures. However, each of them does not have a cryptic cavity. Rather, the detected volumetric spaces look like molecular surface (**Figure S14B and S14C**). In comparison, our Molearn approach results in IRES conformations with cryptic binding cavities with ligand-binding-compatible volume. At this point, our approach is the first report to generate such IRES conformations.

These observations for IRES suggest that Molearn can be extended, as a proof of concept, to larger RNA systems to generate conformations not sampled in the corresponding *apo* MD ensembles. Nonetheless, this extension should be regarded as system-dependent, and the present Molearn model requires further improvement before it can be broadly applied to RNA conformational generation across diverse architectures.

#### SI-13. Future directions for RNA generative modeling toward cryptic-site SBDD

As a conclusion of this subsection, we discuss three technical issues that remain in the current Molearn implementation and outline future directions toward the theoretical prediction of functional RNA conformations. First, the performance of RNA conformation generation varies among the ten independently trained Molearn models, reflecting the stochastic nature of model training. This variability is more profound in the application to IRES than that to TAR. **Table S5** summarizes how the number of IRES conformations passing each filtering step varies across the ten trained models. All ten trained Molearn models yield similar numbers of conformations after the first filtration based on $\%_{\mathrm{TotalErr}}$ (left column in **Table S5**). In contrast, the model indexed as No. **5** accounts for the majority of conformations retained after the second filtration based on cavity pocket size (center column in **Table S5**). This imbalance persists through the third filtration using PoseBusters (right column in **Table S5**). Consequently, most docked IRES–DMA-135 poses ultimately originate from model No. **5**. Ideally, RNA generative models should robustly and consistently explore broad conformational space, regardless of stochastic variations during training.

Second, Molearn-generated conformations do not sufficiently reproduce experimentally derived RNA conformational ensembles at the global-fold level, even when locally ligand-bindable structures are obtained. This limitation manifests as a mismatch between locally ligand-bindable motifs and the global RNA fold. In both TAR and IRES systems, Molearn-generated conformations can form cryptic ligand-binding cavities that are compatible with ligand accommodation, while the overall RNA architecture deviates from experimentally resolved structures. In the IRES case, we selected DMA-135–binding-compatible conformations primarily based on the volume of the cryptic binding cavity, because imposing additional geometrical constraints resulted in no remaining candidate structures. Although geometrical features characterizing the

IRES cryptic binding cavity were identified (**Fig. S14**), none of the Molearn-generated IRES conformations simultaneously satisfied these additional criteria together with the cavity-volume requirement. Consistent with this observation, ligand-docking simulations suggest possible binding of DMA-135 to IRES; however, the detailed local atomistic interaction patterns are not fully reproduced compared with the experimentally resolved complex (**Fig. S13C** versus **Fig. S17B** and **S17C**; *see* also **Table 6**). A similar scale-separation issue is observed in the TAR system. While locally well-arranged structures resembling the Cyt30–Ade35 cryptic binding pocket were obtained, the corresponding global RNA conformations were not faithfully reproduced, with interhelical bent conformations generated instead of the experimentally observed coaxial stacking structures (**Table S7**). Importantly, this scale-separation issue is not specific to a single RNA system but reflects a general limitation of the current Molearn framework. Taken together, these observations indicate that the current generative power of Molearn is not yet sufficient to simultaneously reproduce locally ligand-bindable structures and experimentally accurate global RNA folds, and that this limitation becomes increasingly pronounced for larger and more topologically complex RNA molecules.

Third, the generated conformations may exhibit limited atomistic accuracy, particularly in TAR bulge and loop regions that undergo substantial structural changes between distinct conformational states (*cf.* **Fig. 1A–B**), and therefore require post-generation energetic optimization prior to their use in ligand docking simulations. Such post-generation optimization incurs substantially higher computational cost than Molearn training itself (5 CPU * 8.6 days vs. 500 CPU * 29/24 days, making 14-fold larger computational cost for QRNAS computations than training of Molearn model, in the case of TAR).

Recalling the above three technical issues, we propose to improve Molearn's framework. Addressing these limitations will require deep learning frameworks with greater descriptive power than 1D CNN employed for the current version of Molearn. [15] Use of Diffusion Denoising Probabilistic Model (DDPM) with SE(3) equivariant architecture often leads to remarkable successes in the field of biomacromolecule structure prediction [20,21], so that employing this kind of models could be one possible option for future development of Molearn (or a new deep learning methods) to successfully predict functional RNA conformations. Toward making Molearn a routine application to generate functional RNA structures, we will address the above three issues. The improvement of Molearn could be executed by using established DL frameworks such as DDPM, so that updating Molearn should be well-considered but technically feasible task.

Recent advances in RNA generative modeling [22,23] illustrate how such architectural limitations might be addressed. For example, DynaRNA [22] employs a denoising diffusion probabilistic model with an E(3)-equivariant representation, enabling more faithful preservation of atomic geometry and long-range spatial coherence than one-dimensional convolutional architectures. DynaRNA was designed to capture intrinsic RNA conformational dynamics and has successfully reproduced rarely populated RNA states. Such geometry-aware diffusion models may also reduce the computational cost of downstream structure refinement by generating RNA conformations with higher intrinsic geometric fidelity (*cf.* **Materials and Methods** and Section **SI-11** in Supporting Information). However, it has not been developed to generate or identify ligand-induced cryptic binding conformations, nor to model transition pathways coupled to small-molecule binding. Accordingly, while diffusion-based, geometry-aware frameworks such as DynaRNA offer a promising direction for improving global structural fidelity, extending such approaches to address ligand-induced cryptic site formation would require additional methodological development beyond the scope of the present study. Therefore, studies in this line are left as a future challenge.

### Supporting tables

**Table S1**. Spikiness of RNA BRiQ score-based profile. Unit: $k_B T$.

| Molearn model label | interpolation | pathway search |
|---|---|---|
| 1 | 365.3 | 36.1 |
| 2 | 289.3 | 144.9 |
| 3 | 341.0 | 64.3 |
| 4 | 175.6 | 104.7 |
| 5 | 251.8 | 98.7 |
| 6 | 199.2 | 92.6 |
| 7 | 273.7 | 60.1 |

†Spikiness is defined by difference between the maximum and averaged RNA BRiQ score calculated over all Molearn-generated conformations.

**Table S2**. Statistics of $\%_{TotalErr}$ of Molearn-generated HIV-1 TAR conformations which are mapped inside and outside the convex polygon enclosing MD-derived latent space coordinates. Error estimation denotes 95% confidential interval calculated from standard error. The number in the parentheses denotes grid points in the region.

| Molearn model # | inside | outside |
|---|---|---|
| 1 | 3.81±5.79 (3836) | 8.72±9.63 (6365) |
| 2 | 3.27±4.57 (4417) | 6.96±7.12 (5784) |
| 3 | 1.75±0.58 (5431) | 7.13±7.78 (4770) |
| 4 | 2.53±2.07 (5663) | 7.38±7.46 (4538) |
| 5 | 6.10±7.75 (4374) | 8.01±10.19 (5827) |
| 6 | 3.32±3.80 (3636) | 7.86±7.76 (6565) |
| 7 | 1.88±1.41 (5961) | 7.06±7.25 (4240) |

**Table S3**. Statistics of RNA BRiQ score of Molearn-generated HIV-1 TAR conformations which are mapped inside and outside the convex polygon enclosing MD-derived latent space coordinates. Error estimation denotes 95% confidential interval calculated from standard error. The number in the parentheses denotes grid points in the region. Unit: $k_BT$.

| Molearn model # | inside | outside |
|---|---|---|
| 1 | 245.97±575.82 (3836) | 786.28±1214.74 (6365) |
| 2 | 236.56±602.03 (4417) | 688.62±1143.02 (5784) |
| 3 | −2.90±64.55 (5431) | 526.28±855.73 (4770) |
| 4 | 169.05±147.74 (5663) | 525.11±683.18 (4538) |
| 5 | 472.23±786.61 (4374) | 838.50±1476.99 (5827) |
| 6 | 190.13±357.62 (3636) | 665.86±866.19 (6565) |
| 7 | 17.75±177.09 (5961) | 647.43±1049.95 (4240) |

**Table S4**. Statistics of RNA BRiQ scores for three sets of HIV-1 TAR conformations. Unit: $k_BT$.

| dataset† | mean | median | std. | min. | max. |
|---|---|---|---|---|---|
| MD snapshots | -4.8 | -8.8 | 17.0 | -41.0 | 79.8 |
| model 1 | 583.1 | 58.3 | 1055.5 | -34.7 | 6580.6 |
| model 2 | 492.9 | 77.8 | 973.6 | -9.3 | 8965.4 |
| model 3 | 244.5 | 1.9 | 643.7 | -49.9 | 6912.7 |
| model 4 | 327.4 | 144.2 | 501.1 | 29.8 | 3879.2 |
| model 5 | 496.3 | 81.1 | 761.9 | -13.4 | 5756.2 |
| model 6 | 279.5 | 7.3 | 756.9 | -49.7 | 7398.0 |
| model 7 | 117.9 | 44.3 | 217.3 | -9.4 | 2770.5 |
| model 8 | 62.0 | -9.6 | 303.6 | -50.7 | 4131.1 |
| model 9 | 54.5 | -19.3 | 237.3 | -51.9 | 2347.7 |
| model 10 | 681.5 | 71.1 | 1242.7 | -33.3 | 9075.2 |
| bindable conf. | 200.1 | 162.6 | 109.9 | 23.5 | 551.7 |

†MD snapshots: 400 MD snapshots used for the Molearn training dataset; model No. (Number ranges from 1 to 10): Molearn-generated conformations sampled from 10201 grid points on the two-dimensional latent space. Indexes for models are consistent with those in other Figures, *e.g.*, **Figure S8**; bindable conf.: 61 MV2003-bindable conformations identified in this study.

**Table S5**. Number of Molearn-generated IRES conformations passing filtering processes.

| trained model ID | filtering criteria | | |
|---|---|---|---|
| | $\%_{TotalErr}$ | cavity volume | PoseBusters [†] |
| 1 | 8545 | 223 | 207 (2230) |
| 2 | 8570 | 10 | 20 (100) |
| 3 | 8544 | 20 | 17 (200) |
| 4 | 6926 | 25 | 24 (260) |
| 5 | 9745 | 3439 | 11939 (34390) |
| 6 | 8125 | 28 | 14 (280) |
| 7 | 8662 | 74 | 55 (740) |
| 8 | 8775 | 18 | 32 (180) |
| 9 | 8747 | 38 | 54 (380) |
| 10 | 8203 | 63 | 57 (630) |

[†] Number in parentheses denotes number of conformations tested for PoseBusters, which comes from that of 'cavity volume' multiplied by 10, the number of IRES-bound DMA-135 conformation.

**Table S6.** Local RMSD of Molearn-generated IRES conformations with potential DMA-135-binding cavities relative to NMR-derived DMA-135-bound IRES conformations (PDB entry: 6XB7), calculated using RNA residues A134-A139, U162-A165 (bulge region) . Unit: Å.

| Molearn model # | the number of conformation | ave. with CI95 † | min. RMSd value |
|---|---|---|---|
| 1 | 247 | 7.27±0.04 | 5.38 |
| 2 | 10 | 7.18±0.17 | 5.47 |
| 3 | 20 | 7.26±0.12 | 5.96 |
| 4 | 26 | 7.14±0.11 | 5.54 |
| 5 | 3444 | 6.52±0.01 | 5.38 |
| 6 | 28 | 7.11±0.09 | 5.66 |
| 7 | 79 | 7.37±0.06 | 5.40 |
| 8 | 20 | 7.13±0.11 | 5.34 |
| 9 | 38 | 7.19±0.09 | 5.67 |
| 10 | 63 | 6.91±0.06 | 5.52 |

† For a Molearn generated conformation, each of 10 DMA-135-bound conformations are used to calculate RMSD, then giving 10 values.

**Table S7.** Global RMSD of Molearn-generated IRES conformations with potential DMA-135-binding cavities relative to NMR-derived DMA-135-bound IRES conformations (PDB entry: 6XB7), calculated using all RNA residues. Unit: Å.

| Molearn model # | the number of conformation | ave. with CI95 † | min. RMSd value |
|---|---|---|---|
| 1 | 247 | 10.93±0.06 | 6.99 |
| 2 | 10 | 11.07±0.33 | 7.52 |
| 3 | 20 | 10.77±0.21 | 7.88 |
| 4 | 26 | 10.79±0.18 | 6.78 |
| 5 | 3444 | 10.42±0.01 | 7.71 |
| 6 | 28 | 10.61±0.17 | 7.40 |
| 7 | 79 | 11.41±0.11 | 7.84 |
| 8 | 20 | 10.77±0.22 | 6.29 |
| 9 | 38 | 11.09±0.15 | 7.56 |
| 10 | 63 | 9.91±0.11 | 6.92 |

### Supporting figures

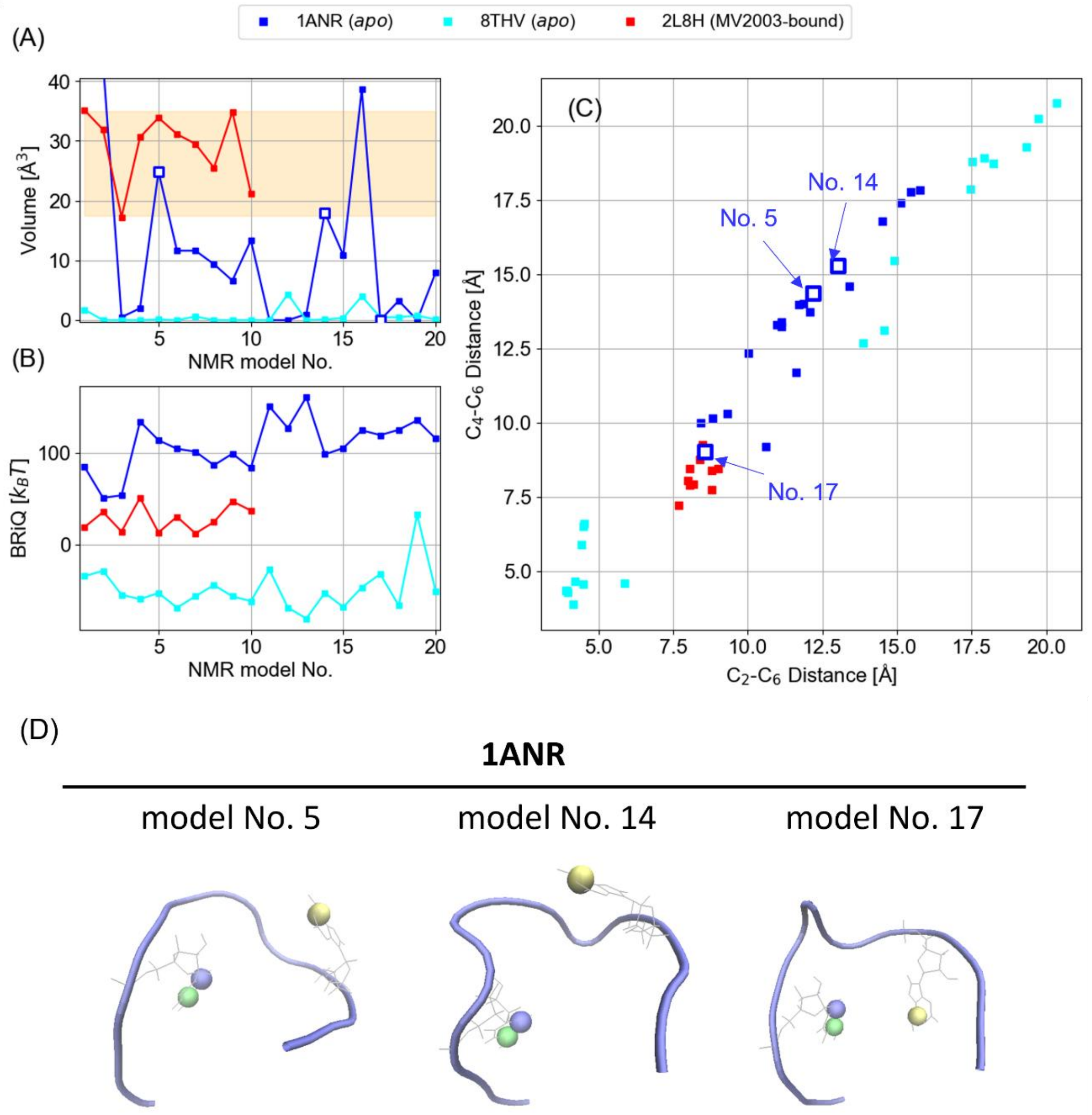


**Figure S1**. Structural characterization of NMR-resolved alternative *apo* HIV-1 TAR conformations (PDB entries of 1ANR and 8THV and). (A) Volume of (potential) MV2003-binding cavity. (B) RNA BRiQ score (BRiQ). (C) Interatomic distances characterizing positional relationship between C30 and A35. (D) Location of C30 and A35 in *apo* form (PDB entry: 1ANR). In panels A and B, orange belts denote value ranges for volume of MV2003-binding cavity and BRiQ[11] for MV2003-bound TAR conformations (PDB entry:

2L8H). In panels A and C, blue open squares are for No. 5, 14 and 17 models of 1ANR. In panel D, blue, green and yellow balls represent C30's $C_2$, C30's $C_4$ and A35's $C_6$ atoms, respectively.

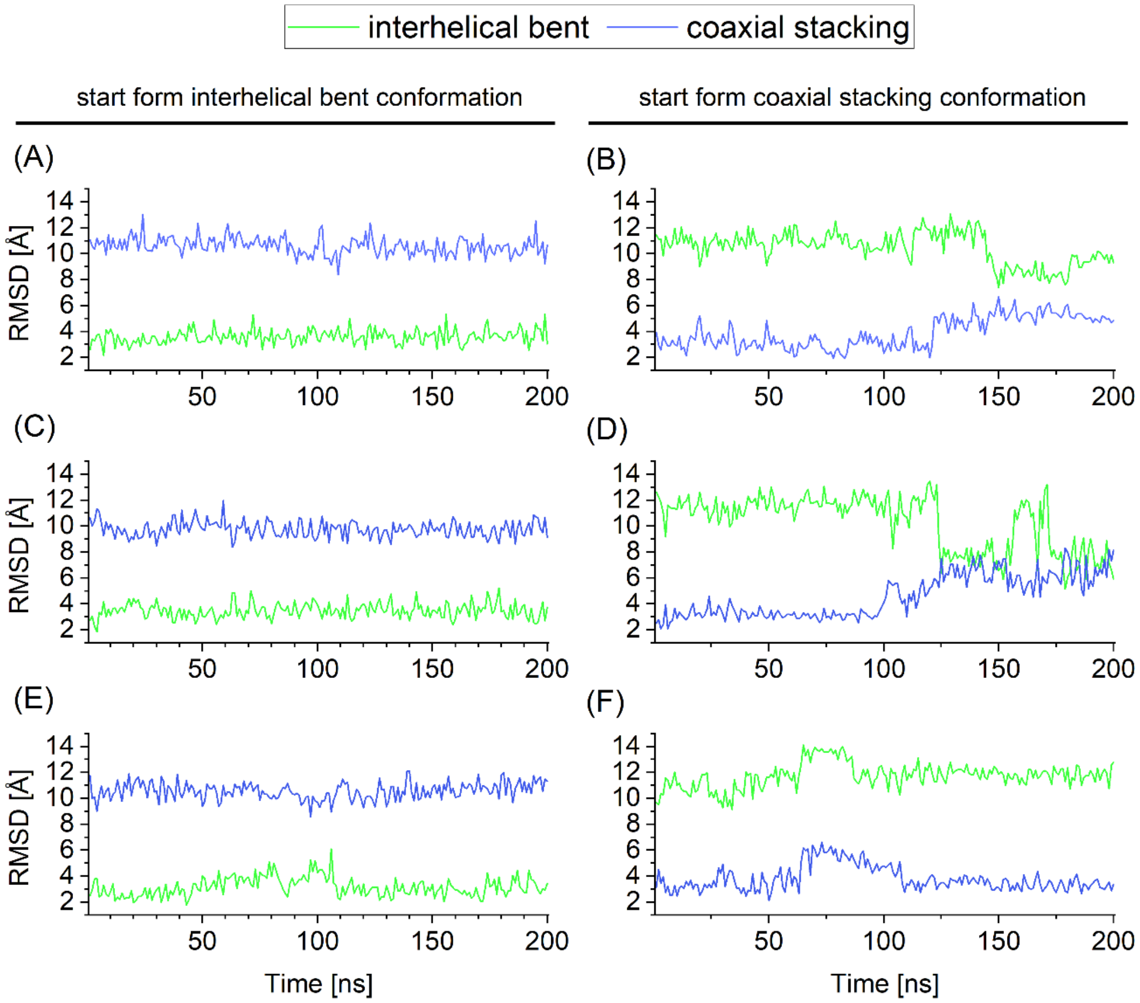


**Figure S2**. Time course analyses of Root Mean Square deviation (RMSD) to NMR-derived representative HIV-1 TAR conformations. (A), (C) and (E) are for 200-ns NPT MD simulations starting from interhelical bent, while (B), (D) and (F) are for those starting from coaxial staking (*see* **Fig. 1B**), respectively. RMSD values to NMR-derived interhelical bent and coaxial staking conformations are shown by green and blue lines, respectively.

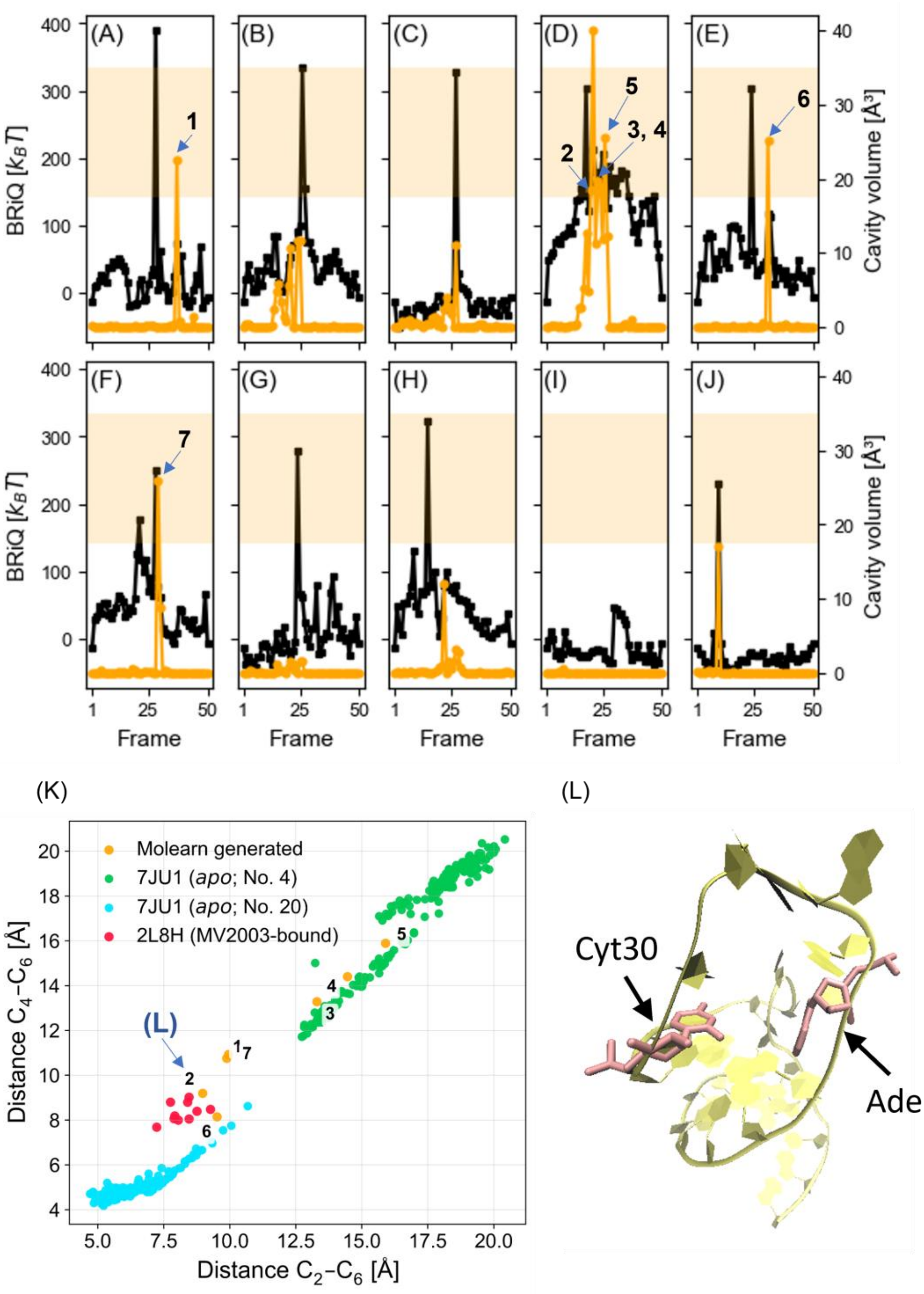


**Figure S3**. Molearn-generated transition path conformations with the 1000th epoch model. Panels A to J correspond to each of the ten Molearn models, numbered from 1 to 10 (*c.f.*, **Fig. 5**). The black and orange lines are for RNA BRiQ score (annotated as BRiQ in panels) [Unit: $k_BT$] and cavity volume [Unit: $Å^3$], respectively. the orange belt denotes the value range for NMR-resolved HIV-1 TAR–MV2003 complexes. Conf. [#] denotes index of TAR conformation on transition pathway: #1 and #50 are for the interhelical bent and coaxial stacking conformations (a pair with the maximum RMSD value, which is picked up from the training dataset and similar to models No. 4 and 20 shown in **Fig 2B**) and the others are for those generated by interpolating the two conformations. The conformations satisfying the volume criterion are annotated by numbers from 1 to 7, which corresponds to those in panel (K). (K) Interatomic distances characterizing positional relationship between C30 and A35. 'L' beside the annotation of **2** denotes the representative conformation shown in the panel L. (L) Representative Molearn-generated TAR numbered by **2**, in panels D and K, satisfying geometrical criteria for cryptic binding formation.

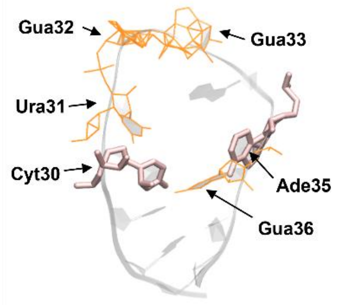


**Figure S4**. Molearn-generated TAR conformation satisfying criteria of cavity volume and interatomic distance pair but failing to perfectly relax steric hindrance, annotated by **6** in **Fig. S3**. RNA backbone is represented by transparent ribbon. Cyt30 and Sde35 are colored by pink. Residues with steric hindrance are illustrated by orange sticks.

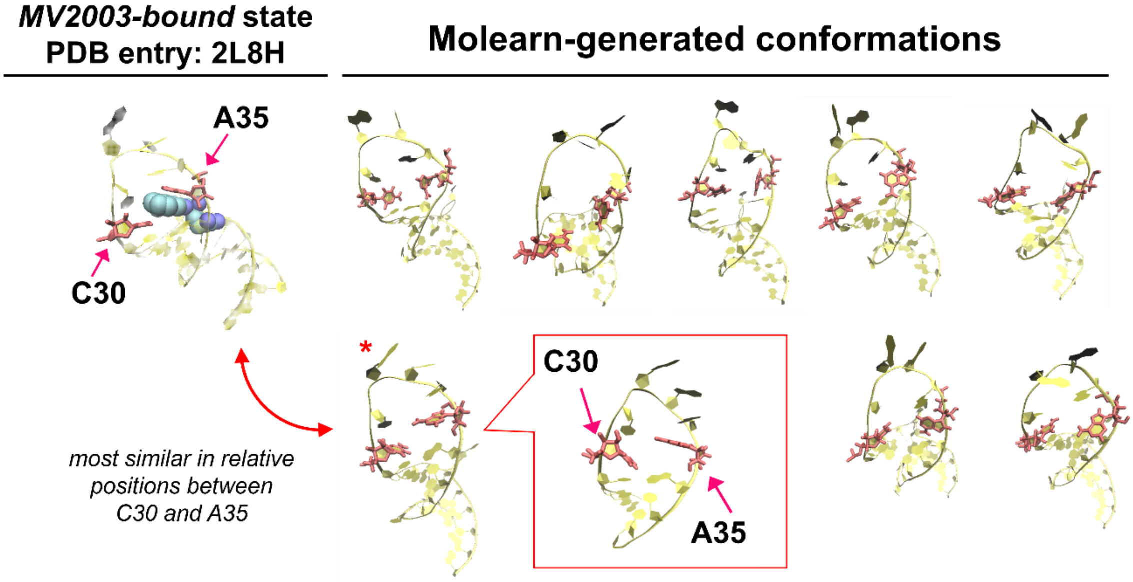


**Figure S5**. Representatives of MV2003-binding-compatible conformations, observed among the 61 conformations generated by Molearn. NMR-derived MV2003-bound HIV-1 TAR is shown at the left, as reference. The Molern-generated conformation annotated by red star is the most similar to the NMR-derived one.

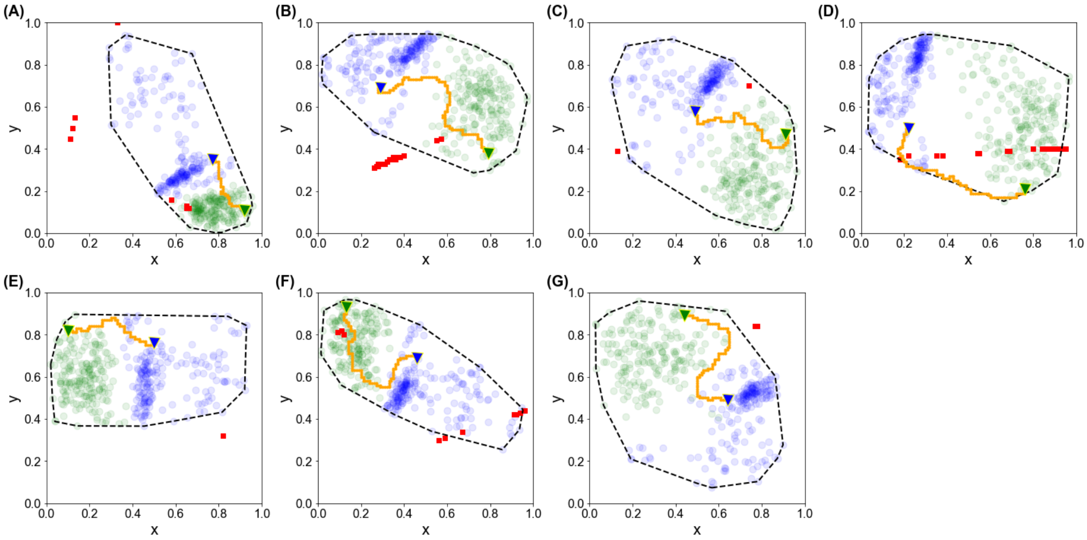


**Figure S6**. Mapping HIV-1 TAR conformations on Molearn's latent space derived from each of seven trained models. (A) to (G) correspond to No. 1 to No. 7 in **Fig. 5**. Red squares denote Molearn generated conformations with potential MV2003-binding cavities. Transparent bule and green circles are snapshot structures obtained from MD simulation for interhelical bent and coaxial stacking conformations (*cf*. **Fig. 1B**), respectively. Blue and green inversed triangle are for a pair of MD snapshots with the maximum RMSD value. Minimum RNA BRiQ score-based pathways connecting the inverse triangles are shown by orange lines. Black dotted line denotes a convex polygon covering MD-derived snapshot structures.

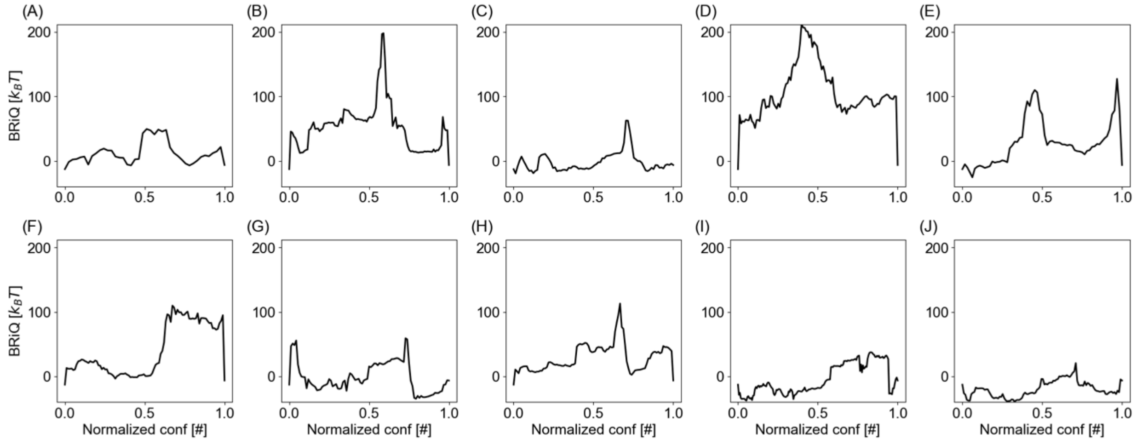


**Figure S7**. RNA BRiQ score (BRiQ) along the minimum score pathway for Molearn-generated conformations at the 1000$^{th}$ training epoch [Unit: $k_BT$]. Panels A to J correspond to each of the ten Molearn models evaluated in **Fig. 5**. Panels A to F are for Molearn models generating HIV-1 TAR conformations with potential MV2003-binding cavity. Since the number of conformations is different among the ten Molearn models, indexes for conformations are normalized between 0 to 1, which referred to as Normalized conf [#]. The values of 0 and 1 are for the representative interhelical bent and coaxial stacking conformations, respectively (a pair with the maximum RMSD value, which is picked up from the training dataset and similar to models No. 4 and 20 in PDB entry: 7JU1, shown in **Fig. 2B**). The others are Molearn-generated conformations on the minimum BRiQ score pathway.

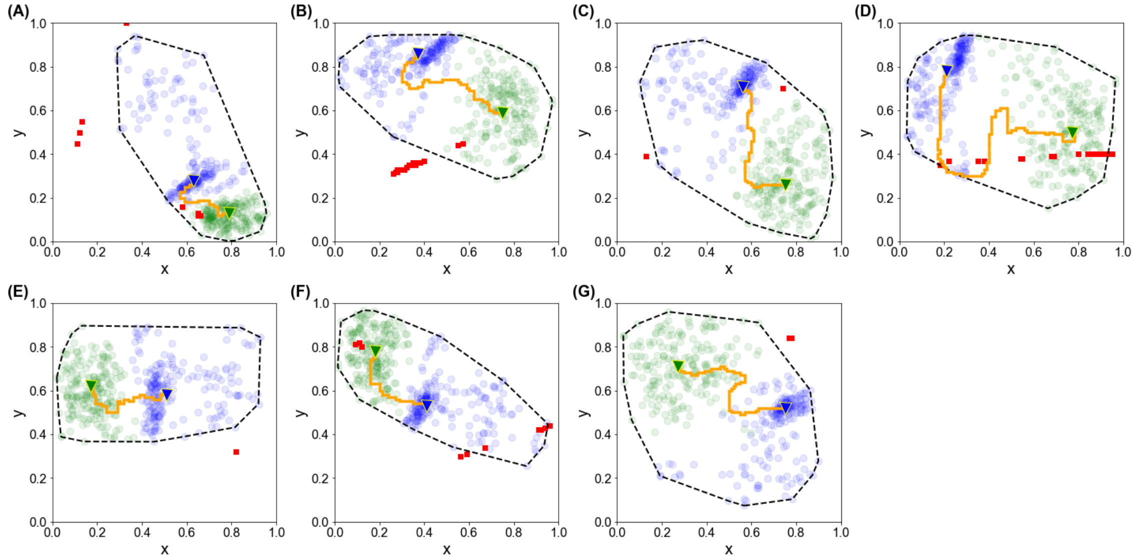


**Figure S8**. Mapping HIV-1 TAR conformations on Molearn's latent space derived from each of six trained models. (A) to (G) correspond to No. 1 to No. 7 in **Fig. 5**. Red squares denote Molearn generated conformations with potential MV2003-binding cavities. Transparent bule and green circles are snapshot structures obtained from MD simulation for interhelical bent and coaxial stacking conformations (*c.f.*, **Fig. 2B**), respectively. Blue and green inversed triangle are for a pair of MD snapshots with the maximum RMSD value. Minimum RNA BRiQ score-based pathways connecting the inverse triangles are shown by orange lines. Black dotted line denotes a convex polygon covering MD-derived snapshot structures.

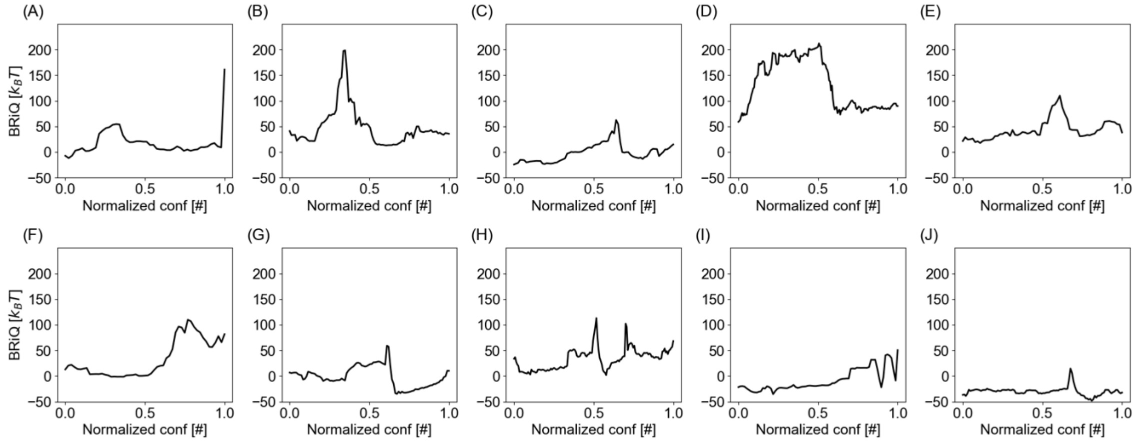


**Figure S9**. RNA BRiQ score (BRiQ) along the minimum score pathway for Molearn-generated conformations at the 1000th training epoch [Unit: $k_BT$]. Panels A to J correspond to each of the ten Molearn models evaluated in **Fig. 5**. Models labeled by A to G generate conformations with potential MV2003-binding cavity. Since the number of conformations is different among the ten Molearn models, indexes for conformations are normalized between 0 to 1, referred to as Normalized conf [#]. Conformations with Normalized conf [#] of 0 and 1 are obtained from the densest regions by Density Based scan clustering (discussed in **SI-6**). The others are Molearn-generated conformations.

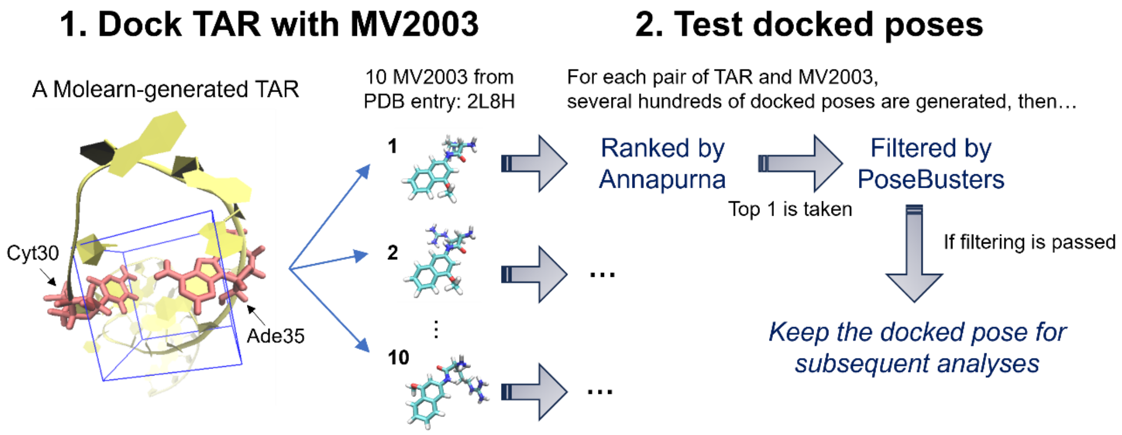


**Figure S10**. Illustration of generation process of HIV-1 TAR–MV2003 docked pose.

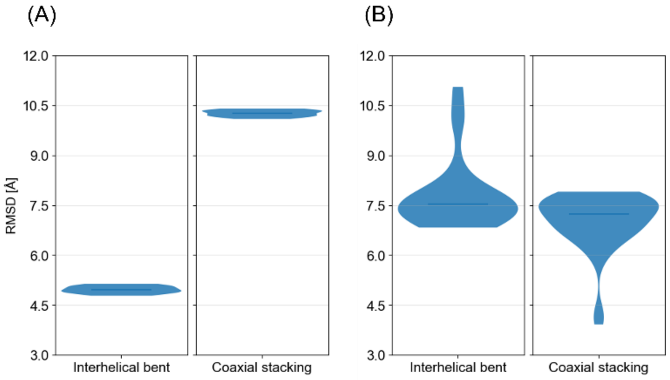


**Figure S11.** Evaluation of global HIV-1 TAR conformation with RMSD to representative *apo* conformations. (A) 10 MV2003-bound conformations (PDB entry: 2L8H). (B) 61 Molearn-generated conformations harboring MV2003-bindable cavities. Representative interhelical bent and coaxial stacking conformations are models 4 and 20 of in the *apo* TAR ensemble (PDB entry: 7JU1; *see* **Fig. 1B**).

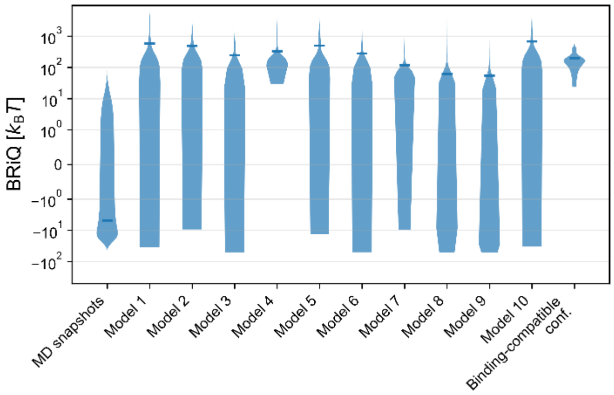


**Figure S12.** Distributions of RNA BRiQ scores for three sets of HIV-1 TAR RNA conformations show with violin plot. The left panel shows the 400 MD snapshots used as the Molearn training dataset. The ten middle panels show Molearn-generated conformations sampled from 10201 grid points on the two-dimensional latent space, with each panel corresponding to an independently trained Molearn model evaluated in **Fig. 5**. The right panel shows the 61 Molearn-generated MV2003-binding-compatible conformations identified in this study.

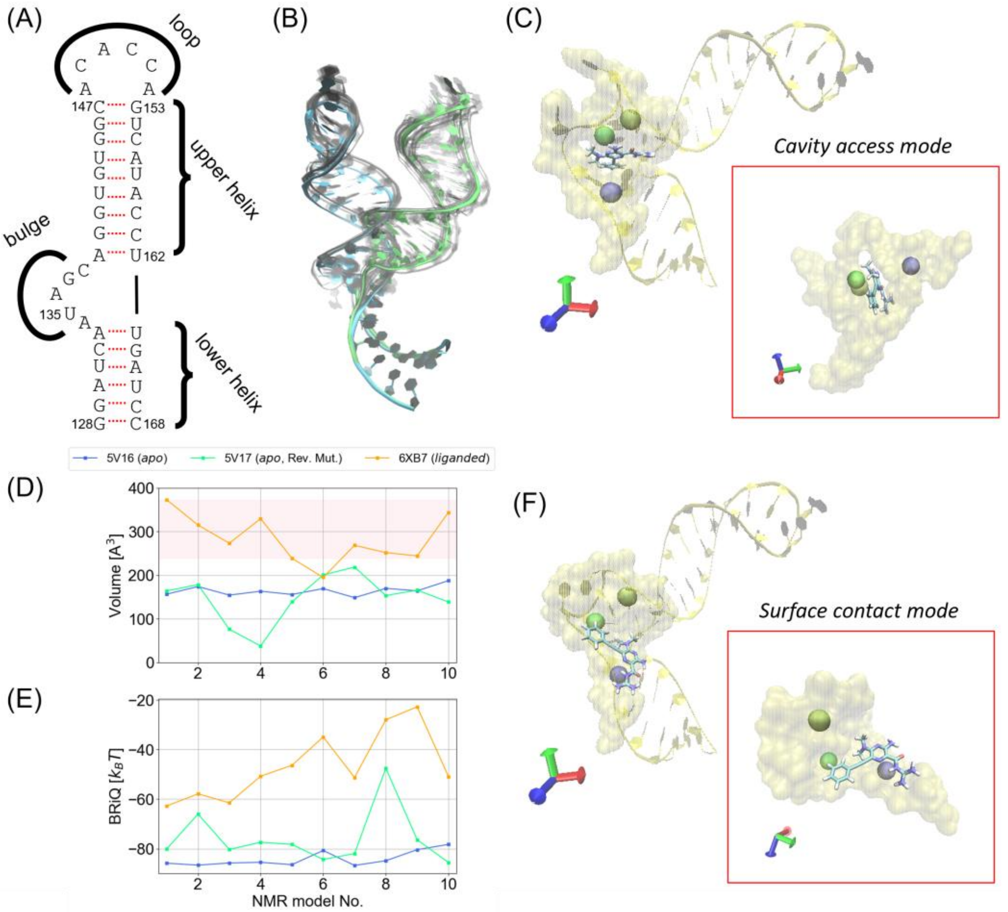


**Figure S13**. Structural and physicochemical characterization of NMR-resolved IRES conformations (*apo*, mutated *apo* and DMA-135 bound forms annotated by PDB entries of 5V16, 5V17 and 6XB7, respectively). (A) Primary and secondary structure of IRES. (B) *apo* IRES structure ensembles. (C) DMA-135 bound IRES structure from model No. 1 of 6XB7, whose binding mode is cavity access. (D) Volume of (potential) DMA-135 binding cavity. (E) RNA BRiQ score (BRiQ). (F) DMA-135 bound IRES structure from model No. 6 of 6XB7, whose binding mode is surface contact. Red dots in panel A denote hydrogen bonding between base pair. In panels B, all of 5V17 models are reverse-mutated (Rev. Mut.) for the physicochemical analyses. RMSD maximum pairs, model No. 10 of 5V17 and model No. 6 of Rev. Mut. 5V17, are colored by blue and green, respectively. The remaining 18 models are shown by gray ribbon. In panel D, orange belts denote value ranges for volume of DMA-135 binding cavity, where DMA-135 accesses IRES's cavity. In panels C and F, IRES and the DMA-135 binding site are shown by yellow ribbon and surface, respectively. blue, green and yellow balls represent C1' atoms of A133, A136 and A142, respectively. Binding of DMA-135 to IRES is highlighted in the red frame.

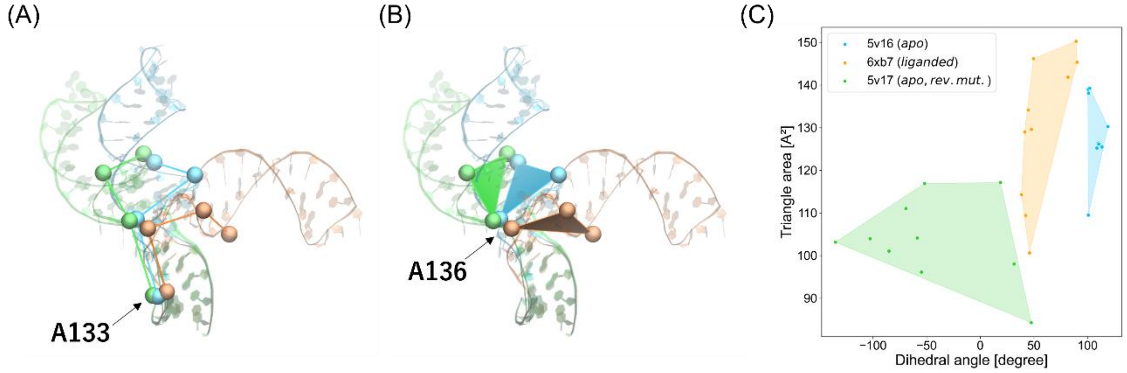


**Figure S14**. Geometry condition discriminating *apo* IRES conformations and those in the DMA-135-bound state. (A) dihedral angle defined by four phosphate atoms derived from A133, A136, A139 and A142. (B) Triangle area defined by three phosphate atoms derived from A136, A139 and A142. (C) Distribution of pair of the dihedral angle and triangle area. In panel C, NMR-resolved conformations are used to calculate geometrical features. Each convex hull covers all IRES conformations for a given PDB entry.

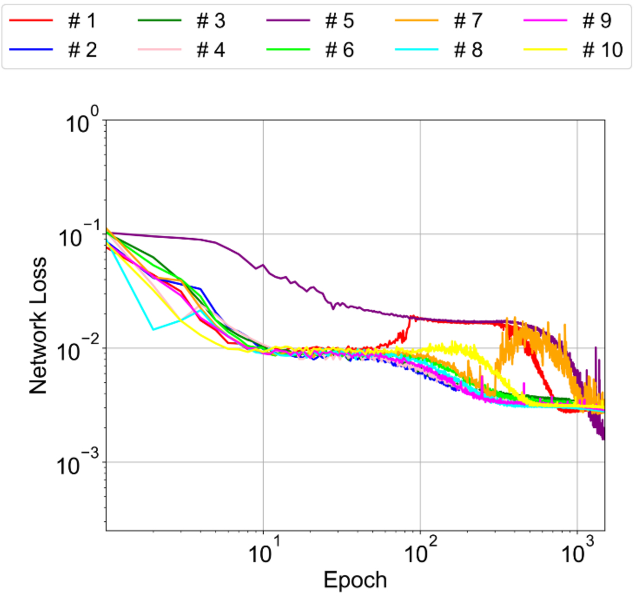


**Figure S15**. Network loss of trained Molean model for IRES. Value at each epoch is an average over 1,000 training steps in the epoch. #1 to #10 in the legend box denote the ten independently trained Molearn models.

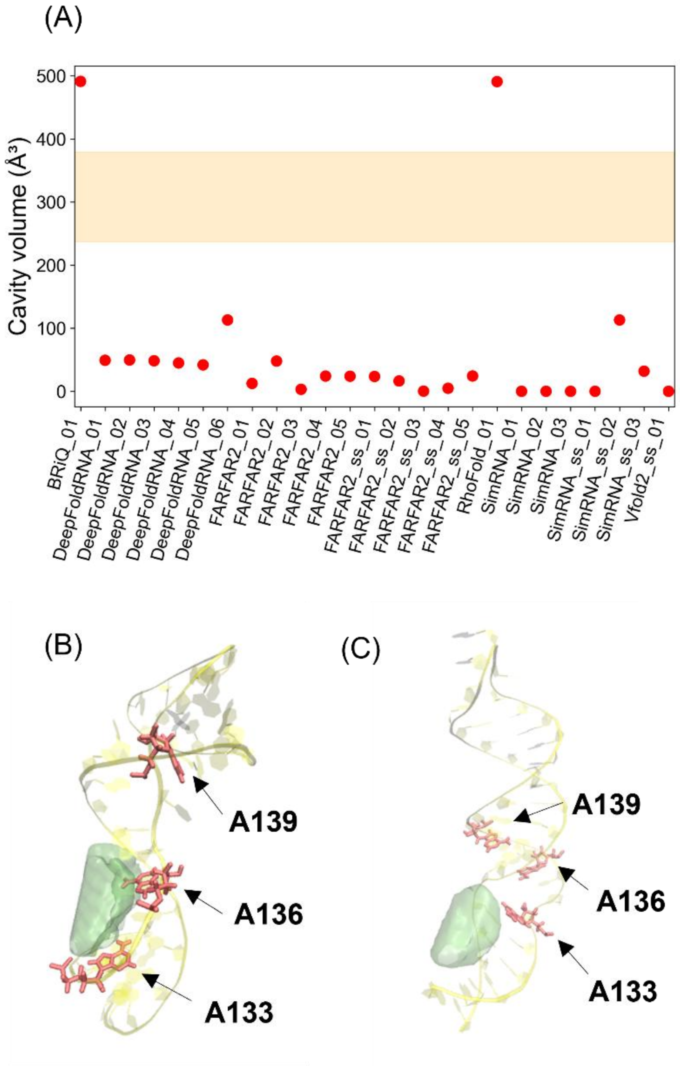


**Figure S16.** Potential DMA-135 binding cavity, calculated by using IRES structures discussed in Nithin's study[24]. Panel (A) shows *c*avity volume. Orange belts denote value ranges for volume of DMA-135 binding cavity calculated by using NMR-derived IRES conformations in the DMA-135-bound state (PDB entry: 6XB7). Structure annotations come from PDB filenames distributed from the authors (https://data.mendeley.com/datasets/8yg88x7rdk/3). Cavity volume calculation failed for Vfold2_ss_02 so that we do not discuss the model. Cavity volume calculation failed for Vfold2_ss_02 so that we do not discuss the model. Panels (B) and (C) are IRES conformations generated by RNA-BRIQ [11] and RhoFold [19], respectively. In these two panels, the ligand and the interacting residues are shown by van der Waals spheres and red sticks, respectively. IRES structure is illustrated by transparent yellow ribbon. Calculated ‘cavity’ is represented by transparent green surface.

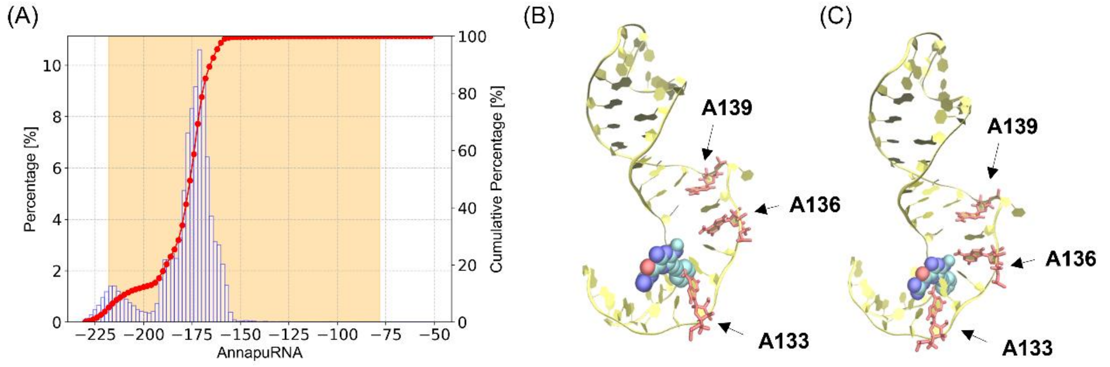


**Figure S17**. Predicted IRES–DMA-135 binding pose. (A) IRES–DMA-135 binding scores calculated by AnnapuRNA scoring function. The orange belt denotes the value range for NMR-resolved IRES–DMA-135 complexes. Blue open bars and red closed circles are for percentages and cumulated percentage of the histogram, respectively. (B) Binding pose with the minimum AnnapuRNA score. (C) Binding pose with the minimum AnnapuRNA score within the orange belt in panel A. In panels B and C, the DMA-135 and the interacting residues are shown by van der Waals spheres and red sticks, respectively. IRES structure is illustrated by yellow ribbon, as in the case of **Fig. S16**.
